# Stable Isotope Tracing in vivo Reveals A Metabolic Bridge Linking the Microbiota to Host Histone Acetylation

**DOI:** 10.1101/2021.07.05.450926

**Authors:** Peder J. Lund, Leah A. Gates, Marylene Leboeuf, Sarah A. Smith, Lillian Chau, Elliot S. Friedman, Mariana Lopes, Yedidya Saiman, Min Soo Kim, Christopher Petucci, C. David Allis, Gary D. Wu, Benjamin A. Garcia

**Affiliations:** Penn Epigenetics Institute, Dept. of Biochemistry and Biophysics, University of Pennsylvania, Philadelphia, PA 19104; Dept. of Medicine, Division of Gastroenterology and Hepatology, University of Pennsylvania, Philadelphia, PA 19104; Laboratory of Chromatin Biology and Epigenetics, The Rockefeller University, New York, NY 10065; Metabolomics Core, Penn Cardiovascular Institute, Perelman School of Medicine, University of Pennsylvania, Philadelphia, PA 19104

**Keywords:** Host-microbiota interactions, epigenetics, histone acetylation, fatty acid metabolism, colitis

## Abstract

The gut microbiota influences host epigenetics by fermenting dietary fiber into butyrate. Although butyrate could promote histone acetylation by inhibiting histone deacetylases, it may also undergo oxidation to acetyl-CoA, a necessary cofactor for histone acetyltransferases. Here, we find that epithelial cells from germ-free mice harbor a loss of histone H4 acetylation across the genome except at promoter regions. Using stable isotope tracing *in vivo* with ^13^C-labeled fiber, we demonstrate that the microbiota supplies carbon for histone acetylation. Subsequent metabolomic profiling revealed hundreds of labeled molecules and supported a microbial contribution to host fatty acid metabolism, which declined in response to colitis and correlated with reduced expression of genes involved in fatty acid oxidation. These results illuminate the flow of carbon from the diet to the host via the microbiota, disruptions to which may affect energy homeostasis in the distal gut and contribute to the development of colitis.

## INTRODUCTION

The distal gut is inhabited by the microbiota, an enormous community of microbes that forms a dynamic ecosystem with its host organism. Interactions within and across kingdoms have a significant impact on host physiology. For instance, the microbiota has roles in preventing enteric infections (Ghosh *et al*, 2011; Ferreyra *et al*, 2014; Ng *et al*, 2013; Baümler & Sperandio, 2016), regulating the immune system (Smith *et al*, 2013; Honda & Littman, 2016; Thaiss *et al*, 2016; Rooks & Garrett, 2016; Gensollen *et al*, 2016) and intestinal epithelium (Rakoff-Nahoum *et al*, 2004; Sanos *et al*, 2009; Levy *et al*, 2015; Kelly *et al*, 2015; Kaiko *et al*, 2016), and assisting with nutrient acquisition (Blanton *et al*, 2016; Sonnenburg & Bäckhed, 2016). Undesirable alterations in the microbiota lead to a state known as dysbiosis (Tiffany & Bäumler, 2019; Gillis *et al*, 2018; Byndloss *et al*, 2017; Lupp *et al*, 2007; Ferreyra *et al*, 2014; Winter *et al*, 2013; Sonnenburg & Bäckhed, 2016; Sonnenburg & Sonnenburg, 2014). Characterized by a loss of diversity and an outgrowth of the facultatively anaerobic Proteobacteria over the obligate anaerobes that typically predominate, dysbiosis is a hallmark of several human diseases, especially ulcerative colitis and Crohn’s disease (Ott *et al*, 2004; Manichanh *et al*, 2006; Frank *et al*, 2007; Ni *et al*, 2017). This correlation motivates the study of host-microbiota interactions in pursuit of understanding the processes that maintain gut homeostasis and how these processes become corrupted in disease.

Host-microbiota interactions depend in part on small molecule metabolites that can function as receptor ligands, enzyme inhibitors, or metabolic precursors (Levy *et al*, 2016; Arpaia *et al*, 2013; Cohen *et al*, 2017; Miyamoto *et al*, 2015; Klepsch *et al*, 2019; Melhem *et al*, 2019). This microbial metabolome is related to the composition of the microbiota and host diet (Marcobal *et al*, 2013; Li *et al*, 2008; Schroeder & Bäckhed, 2016; Martin *et al*, 2007; Walker *et al*, 2014). Fermentation of dietary fiber by the microbiota generates high concentrations of the short-chain fatty acids (SCFAs) acetate, propionate, and butyrate (Ríos-Covián *et al*, 2016; Koh *et al*, 2016; Morrison & Preston, 2016). Butyrate has received much attention because of its ability to inhibit histone deacetylases (HDACs) (Candido *et al*, 1978). HDACs represent one of the many classes of chromatin modifying enzymes that modulate the post-translational modification status of histone proteins, thereby contributing to the epigenetic regulation of gene expression (Allis & Jenuwein, 2016). Histones, which form octameric scaffolds that associate with 147 bp of DNA to organize eukaryotic chromatin into nucleosomes, are subject to a myriad of post-translational modifications, among which lysine acetylation and methylation are the most appreciated and well-studied (Fan *et al*, 2015; Kouzarides, 2007; Berger, 2007; McGinty & Tan, 2015; Zhao & Garcia, 2015). These modifications, added and removed by “writer” and “eraser” enzymes, regulate chromatin activity in part by recruiting “reader” proteins to promote a biological outcome, such as transcriptional activation or repression of the genes encoded in the underlying DNA. Thus, inhibition of HDACs by butyrate has the potential to impact gene expression. Previous work has shown that butyrate, produced primarily by obligate anaerobes, enforces immunological tolerance in the gut through acetylation of histone H3 near the *Foxp3* gene, thereby supporting the differentiation of regulatory T cells (Tregs) (Furusawa *et al*, 2013).

Beyond acting as an HDAC inhibitor, butyrate also functions as a receptor ligand and an energy source. Butyrate and other SFCAs bind to numerous GPCRs (Brown *et al*, 2002). The ameliorative effect of a high-fiber diet on experimentally induced colitis has been linked to GPR43 and GPR109A (Macia *et al*, 2015). Butyrate may also activate intracellular receptors, such as PPARγ (Byndloss *et al*, 2017; Alex *et al*, 2013). Microbiota-dependent PPARγ signaling in epithelial cells promotes mitochondrial β-oxidation, which is essential for maintaining anaerobic conditions and homeostasis in the gut lumen. This metabolism of gut epithelial cells depends heavily on microbial SCFAs, which serve as readily available carbon sources that can undergo β-oxidation to generate acetyl-CoA, thereby fueling the TCA cycle and ATP production (Donohoe *et al*, 2012, 2011). Since acetyl-CoA is the donor substrate for histone acetyltransferases (HATs), butyrate also has the potential to impact histone acetylation patterns through its metabolism (Donohoe *et al*, 2012). As proposed by Donohoe et al. (2012), whether butyrate acts as an indirect activator of HATs or a direct inhibitor of HDACs likely depends on the propensity of cells for fatty acid metabolism, which is expected to be high in the case of gut epithelial cells.

Stable isotope labeling presents an attractive approach to study the contribution of different substrates to downstream metabolic pathways and protein modifications. Using this method, carbon from the medium-chain fatty acid octanoate was recently traced to the acetyl groups of histones (McDonnell *et al*, 2016). Although early studies with radioactive isotopes were instrumental in establishing the importance of butyrate metabolism in the distal gut (Roediger, 1980a, 1980b; Ahmad *et al*, 2000; Chapman *et al*, 1994), their measurement of ^14^CO_2_ only accounts for the contribution of butyrate oxidation to the TCA cycle without consideration of any alternative fates of butyrate-derived acetyl-CoA, such as its use for histone acetylation. Since acetyl-CoA occupies a central hub in cell metabolism, the microbiota has the potential to support many additional metabolic pathways in gut epithelial cells. Accordingly, subsequent studies provided evidence of radioisotope incorporation into lipids and histones, though the identities of these labeled molecules were not further resolved (Andriamihaja *et al*, 2009; Leschelle *et al*, 2000). A more recent investigation with stable isotopes, in which *E. coli* was labeled externally with ^13^C-glucose and then administered to uncolonized mice, demonstrated the remarkable extent to which microbially-derived compounds enter host tissues (Uchimura *et al*, 2018). However, there is a need for alternative strategies to more fully model the metabolic transformations and ecological interactions that occur in an intact microbiota concentrated in the distal gut with nourishment from the host diet.

In the present study, we aimed to broadly investigate the influence of the microbiota on histone acetylation patterns in the distal gut and provide direct evidence of a metabolic connection between the microbiota and host epithelial cells that feeds into histone acetylation. Through deep profiling of histone modifications with mass spectrometry and subsequent ChIP-seq analysis, we find that germ-free mice have a genome-wide loss of histone H4 acetylation except near transcription start sites. Supporting a metabolic contribution of the microbiota to histone acetylation, we show through isotope tracing experiments that histone acetyl groups contain carbon derived from butyrate and dietary fiber. Finally, we combine our novel *in vivo* isotope tracing approach with untargeted metabolomics and complementary high-throughput analyses in a mouse model of intestinal inflammation, revealing how inflammation disrupts integration between the microbiota and host fatty acid metabolism. Overall, these findings highlight how microbial metabolism of dietary fiber contributes to host cell metabolism and histone acetylation in the distal gut.

## RESULTS

### Acetylation of histone H4 across gene bodies is reduced in the absence of the microbiota

To assess the impact of the microbiota on histone modifications in the distal gut, we prepared histone extracts from cecal and colonic epithelial cells isolated from either conventional (Conv) or germ-free (GF) mice and analyzed them by bottom-up mass spectrometry (MS), an approach which is capable of detecting hundreds of uniquely modified histone peptides (Fig. 1A). Overall, we identified 47 histone peptides with statistically significant differences (Supplemental Fig. 1A, 1B). Compared to their colonized counterparts, mice lacking a microbiota had notable changes in the relative abundances of the modified forms of a peptide derived from the N-terminal tail of histone H4 (Fig. 1B, Supplemental Fig. 1A, 1B). This peptide (^4^GKGGKGLGKGGAKR^17^) contains four acetylation sites at H4K4, H4K8, H4K12, and H4K16. Overall levels of mono-, di-, and tri-acetylation on this peptide were significantly reduced in the ceca and colons of GF mice with a corresponding increase in the levels of the unmodified peptide from roughly 50% to over 60% (Fig. 1B). Thus, the microbiota supports histone H4 acetylation in epithelial cells from the cecum and colon, consistent with a previous study that examined whole colon tissue (Krautkramer *et al*, 2016).

**Figure 1.**
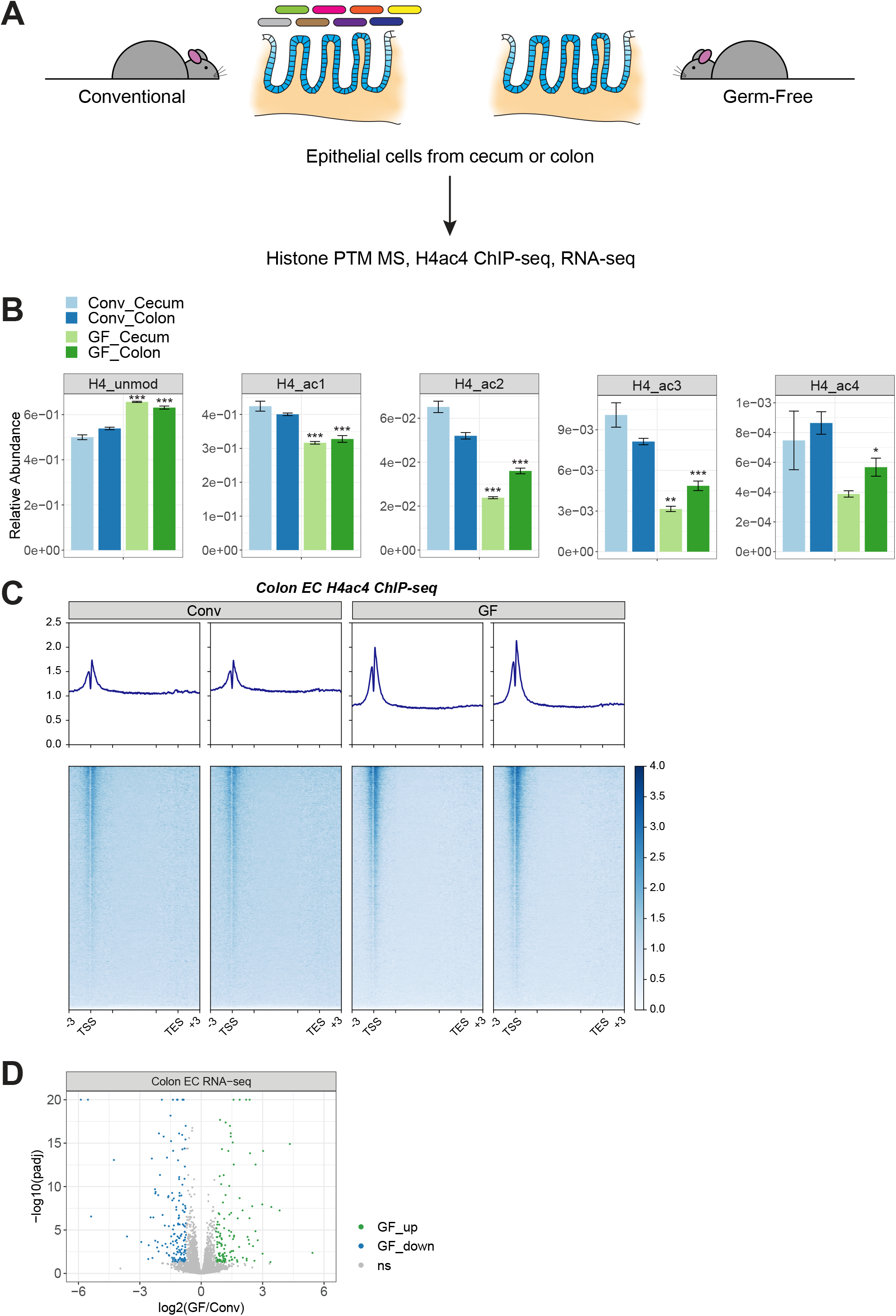
Acetylation of histone H4 across gene bodies is reduced in the absence of the microbiota. A) Schematic of experimental design. Epithelial cells were isolated from the colons and ceca of conventional (Conv) or germ-free (GF) mice. Histone post-translational modifications were profiled by mass spectrometry. Acetylated H4 was profiled in colonic epithelial cells by ChIP-seq in parallel with gene expression profiling by RNA-seq. B) The relative abundances of the unmodified and acetylated forms of the histone H4 tail peptide (^4^GKGGKGLGKGGAKR^17^) are plotted for epithelial cells from ceca or colons of GF versus Conv mice (mean ± sem, n = 5). The abundances of the mono-, di-, and tri-acetylated forms (ac1, ac2, ac3), were obtained by summing the relative abundances of all peptides carrying a given number of acetyl groups regardless of the specific positions of the acetyl groups. * p < 0.05, ** p < 0.01, *** p < 0.001 by unpaired, two-tailed t-test comparing germ-free and conventional for a given tissue. C) Heatmaps and averaged profile plots of the reference-normalized H4ac4 ChIP signal across all genes for two biological replicates from conventional or germ-free mice. Profile plots span from 3 kb upstream of the transcription start site (TSS) to 3 kb downstream of the transcription end site (TES). The distance between the TSS and TES is scaled to a uniform length except for 3 kb downstream of the TSS and upstream of the TES, denoted by the internal tick marks. D) Volcano plot of fold-change versus statistical significance for the expression of 12,094 genes in colonic epithelial cells from conventional and germ-free mice. Significant differences (q < 0.05, log2 FC > mean ± 2 × sd) are highlighted in green and blue. Adj. p-values exceeding 1e-20 are plotted at 1e-20. n = 5. ns: not significant.

To augment our analysis of the overall levels of different histone modifications, we next performed ChIP-seq to localize the reduction in H4 acetylation to specific genomic regions. Given that global alterations in histone marks may obscure the detection of quantitative changes by standard ChIP-seq approaches, we included a spike-in of exogenous chromatin from *Drosophila* S2 cells to serve as an external reference (Orlando *et al*, 2014). Consistent with the reduction in H4 acetylation observed by mass spectrometry, the proportion of sequencing reads originating from the exogenous chromatin was higher in the GF samples, indicating less enrichment of mouse chromatin (Supplemental Fig. 2A). We observed a modest but consistent reduction in H4 acetylation at gene bodies across the genome rather than a more significant loss at specific loci in GF mice (Fig. 1C). Transcription start sites (TSS) were insulated from this loss and appeared to become even more prominently marked by H4 acetylation. Despite the overall decrease in gene body acetylation, GF mice had relatively small and balanced numbers of 125 upregulated and 165 downregulated genes (Fig. 1D, Table 1), similar to previous work (Camp *et al*, 2014). Pathway analysis showed an enrichment for small molecule and ion transporter activity, oxidoreductase activity, and brush border and extracellular matrix components among these differentially expressed genes (Supplemental Fig. 2B).

### Acetylated histones contain carbon derived from butyrate and dietary fiber

A lack of microbiota-dependent butyrate, and therefore a release on the inhibition of HDACs, could explain the reduced levels of H4 acetylation in GF mice. However, in addition to its activity as an HDAC inhibitor, butyrate constitutes a major energy source for gut epithelial cells via its oxidation to acetyl-CoA, the donor substrate for HATs (Donohoe *et al*, 2011, 2012). Thus, a lack of butyrate may also decrease acetyl-CoA levels and HAT activity, leading to a reduction in histone H4 acetylation in GF mice. To determine whether butyrate contributes to histone acetylation through metabolism, we incubated Caco2 cells with 1 mM ^13^C-labeled butyrate and then assayed isotope incorporation into acetylated histone peptides by MS (Fig. 2A). As depicted in Fig. 2B, the isotope label appeared on acetylated histones within 30 mins, reaching a level that then plateaued over 24 hrs. Mass spectra of precursor ions showed a shifted isotopomer distribution with increased isotopomer intensities beginning with the m+2 peak, consistent with the presence of acetyl groups composed of two ^13^C atoms (Supplemental Fig. 3A). Isotope incorporation ranged from less than 1% to almost 40% and seemed to be dictated mainly by the total number of acetyl groups on the peptide (Fig. 2B). The multiply acetylated forms (ac2, ac3, ac4) of the histone H4 tail peptide attained higher levels of labeling than its mono-acetylated forms (ac1), possibly because the former is more likely to originate from more accessible regions of chromatin with greater HAT activity. Starving cells of alternative carbon sources, such as glucose, potentiated the isotopic incorporation from butyrate (Supplemental Fig. 3B).

**Figure 2.**
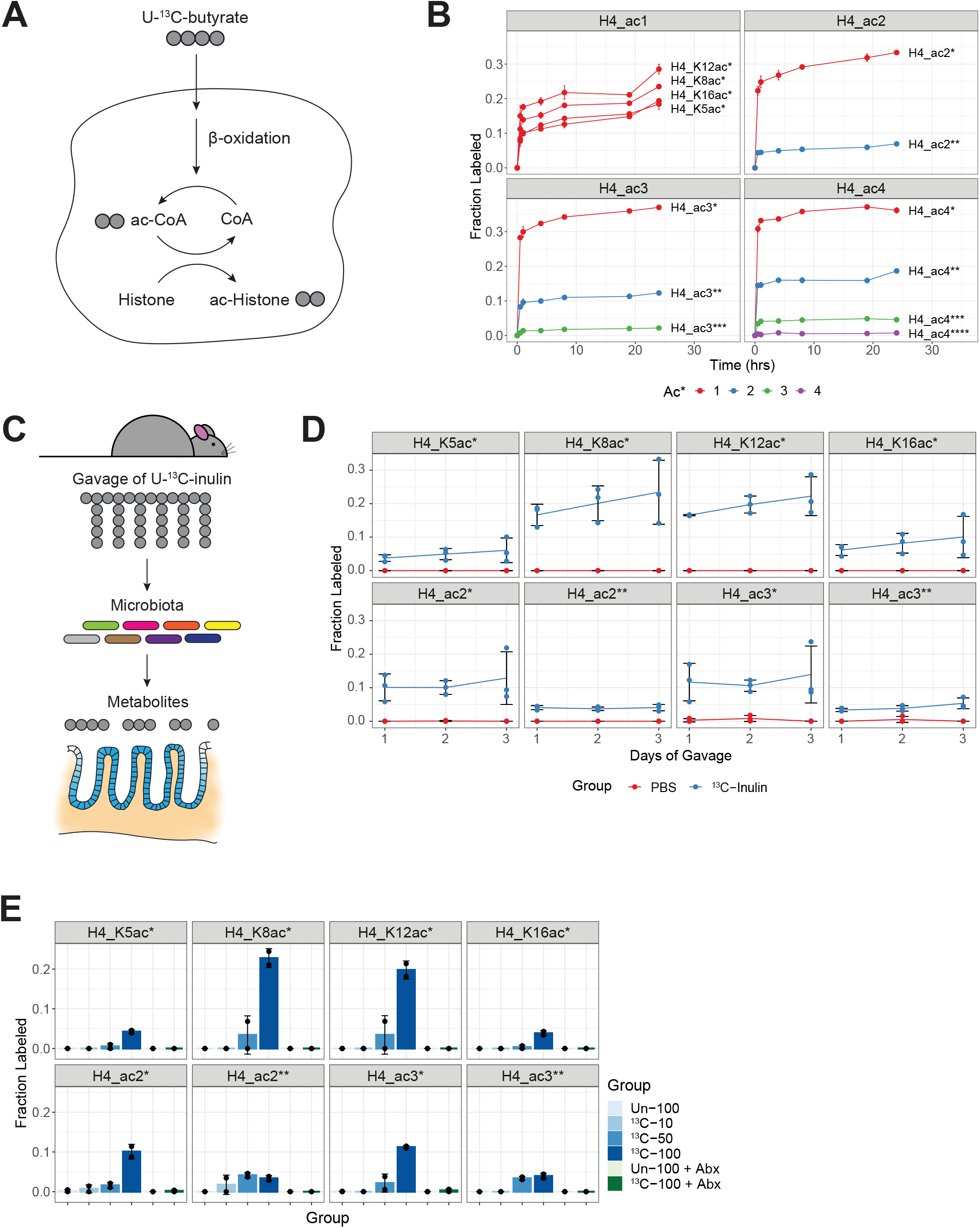
Acetylated histones contain carbon derived from butyrate and dietary fiber. A) Outline of isotope tracing approach with U-^13^C-butyrate in cell culture. B) Caco2 cells were incubated with 1 mM U-^13^C-butyrate over time. Isotope incorporation into histone acetyl groups was determined by mass spectrometry (mean ± sd, n =2-4). The mono-, di-, tri-, and tetra-acetylated forms (ac1, ac2, ac3, ac4) of the histone H4 tail peptide are plotted in separate windows. Within each window, the different combinations of labeled and unlabeled acetyl groups are plotted in different colors. The number of labeled acetyl groups is denoted by asterisks (e.g. ac4** indicates 4 total acetyl groups, of which 2 are ^13^C-labeled). The 0 hr timepoint represents control cells incubated with 1 mM unlabeled butyrate for 24 hrs. C) Outline of isotope tracing approach with U-^13^C-inulin in mice. D) Mice were orally gavaged with ^13^C-inulin or PBS once per day for up to 3 days. Histones were extracted from colonic epithelial cells harvested 24 hrs after the last gavage. Isotope incorporation is plotted over time for various forms of the acetylated histone H4 tail peptide (mean ± sd, n = 3). E) Mice received two daily gavages of labeled (^13^C) or unlabeled (Un) inulin with or without antibiotic (Abx) treatment. Isotope incorporation for the acetylated histone H4 tail peptide is plotted across the different treatment groups (mean ± sd, n = 2). The dosage of inulin (mg) is indicated by numbers.

To test whether butyrate serves as a carbon source for histone acetylation in a more physiological model, we treated mice with isotope-labeled, fermentable fiber (U-^13^C-inulin). Inulin is known to be metabolized into short-chain fatty acids (SFCAs), including butyrate (Deroover *et al*, 2017; Butts *et al*, 2016). Thus, we predicted that the microbiota would ferment inulin into butyrate, which would then be oxidized by gut epithelial cells into acetyl-CoA for use by HATs (Fig. 2C). In line with this hypothesis, we detected isotope incorporation into acetylated histone H4 as well as histone H3 after gavaging mice with ^13^C-inulin but not vehicle control or unlabeled inulin (Fig. 2D and Supplemental Fig. 3C, 3E). Incorporation ranged from 5% to 10% and occurred in a dose- and time-dependent manner (Fig. 2D, 2E and Supplemental Fig. 3C-F). Demonstrating a dependence on the microbiota, antibiotics suppressed isotope incorporation (Fig. 2E and Supplemental Fig. 3D, 3F). These results provide evidence that butyrate and other microbiota-dependent products generated from dietary fiber can act as carbon sources for histone acetylation in the colon.

### Inflammation disrupts the composition, activity, and compartmentalization of the gut microbiota and its associated molecules

Next, we endeavored to trace carbon flow through the intermediary molecules connecting fiber fermentation by the microbiota to acetyl-CoA metabolism and histone acetylation in host cells. Additionally, since inflammatory bowel diseases are commonly associated with alterations in cellular metabolism and the microbiota (Ott *et al*, 2004; Manichanh *et al*, 2006; Frank *et al*, 2007; Roediger, 1980b, 1980a; Ahmad *et al*, 2000; Chapman *et al*, 1994; Haberman *et al*, 2019), we also aimed to determine whether inflammation has a negative impact on carbon flux from the microbiota to gut epithelial cells, which could more broadly affect the metabolic pathways upstream of histone acetylation. To this end, we performed our *in vivo* isotope tracing approach in conjunction with DSS-induced colitis and turned our attention from histones to small molecule metabolites (Fig. 3A). We also carried out additional high-throughput analyses in parallel to gain more insight into the factors, such as host gene expression, that could influence the gut metabolome and carbon transfer.

As expected, mice treated with DSS developed colitis as evident from weight loss (Fig. 3B). We then profiled tissue extracts of healthy and diseased mice with an LC-MS-based platform for untargeted metabolomics, allowing measurement of overall metabolite levels as well as the incorporation of ^13^C atoms. Focusing first on general metabolite levels, we found that each tissue had a distinct metabolome in the healthy state, though the colon and cecum were closely related (Fig. 3C, Table 2). The cecal contents were most disparate from the other tissues, which is not surprising given the capacity of the microbiota to perform metabolic transformations beyond that encoded in the host genome and which are kept compartmentalized by the epithelial barrier. The effect of this barrier is especially apparent by comparing metabolite features in the cecal contents versus other tissues in healthy mice, which display trends of almost complete restriction to, or exclusion from, the lumen (Fig. 3D, Supplemental Fig. 4A). DSS had the greatest impact on the metabolome of the cecal contents, causing a shift in the direction of the colon and cecum (Fig. 3C). This change is likely driven by a compromised epithelial barrier, allowing leakage of host molecules into the lumen, or impairment in the normal output of the microbiota due to disease-associated alterations in its activity or composition. Epitomizing the latter case, deoxycholic acid, hydroxyindoleacetic acid, and urobilinogen, all of which are known to be dependent on the microbiota (Hamoud *et al*, 2018; Kashyap *et al*, 2013; Grüner & Mattner, 2021), declined significantly in the cecal contents of diseased mice (Supplemental Fig. 4A). Conversely, lactic acid and palmitoylcarnitine, which are more associated with host metabolism, became significantly elevated in the cecal contents of diseased mice (Supplemental Fig. 4A). We observed a small number of features showing upregulation in diseased cecal contents beyond the levels seen in healthy tissues, possibly representing colitis-induced molecules from the microbiota or molecules modified by the inflammatory response. These features included caprolactam, dipropylnitrosamine, nitrotyrosine, and ornithine (Supplemental Fig. 4A).

**Figure 3.**
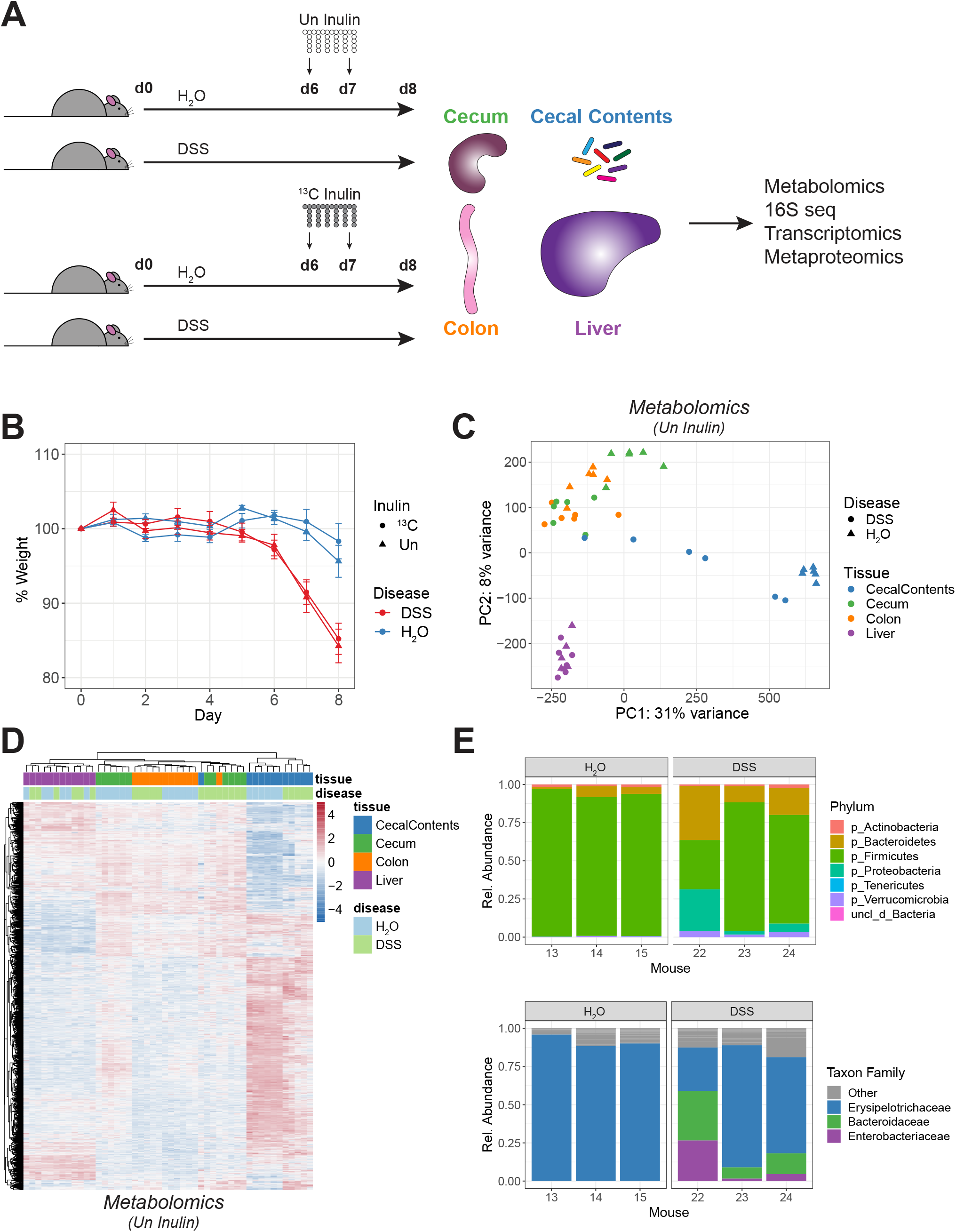
Inflammation disrupts the composition, activity, and compartmentalization of the gut microbiota and its associated molecules. A) Outline of isotope tracing approach in colitis model. Mice were treated with or without DSS to induce colitis. On days 6 and 7, ^13^C-labeled or unlabeled inulin was delivered by gavage. Tissues were harvested on day 8 for analysis. B) Change in relative weight over time. Mean ± sem, n = 6. C) Overall metabolite levels were analyzed in mice receiving Un inulin. PCA was performed using 12,505 features from untargeted metabolomics. D) Heatmap of metabolite features showing differential abundance across tissues or disease (3345 features, q < 0.05 by 2-way ANOVA). E) Relative abundances of different phyla based on 16S sequencing of cecal contents from 3 mice per group (top). Relative abundances of the top 3 OTUs and their family assignments (bottom).

The gut metabolome is determined in part by the composition of the microbiota, which is known to become altered in colitis. Thus, we also performed 16S rRNA sequencing on a subset of samples to assess community composition in healthy and diseased mice and correlate this information with the metabolomics data. In healthy mice treated with ^13^C-inulin, a single OTU accounted for roughly 90% of the entire community (Fig. 3E, Table 3). This unexpected finding contrasts with the high levels of diversity associated with a typical microbiota and suggests that inulin may act as a prebiotic to promote the expansion of this OTU, which matched perfectly to the 16S rRNA of *Dubosiella newyorkensis* (strain NYU-BL-A4) from the *Erysipelotrichaceae* family of the Firmicutes phylum (Cox *et al*, 2017). Inulin has been previously associated with the growth of *Erysipelotrichaceae* (Catry *et al*, 2018; Li *et al*, 2020). Although this OTU persisted at high levels in diseased mice, colitis stimulated dramatic increases in organisms from the Bacteroidetes (OTU2, OTU8), Proteobacteria (OTU3), and Verrucomicrobia (OTU6) phyla (Fig. 3E). The outgrowth of Proteobacteria is a common occurrence in patients with inflammatory bowel disease (Ott *et al*, 2004; Manichanh *et al*, 2006; Frank *et al*, 2007). Not surprisingly, Proteobacteria was associated with more severe colitis based on weight loss (Supplemental Fig. 4B). Conversely, *Erysipelotrichaceae* was positively associated with health.

To complement the 16S data, we characterized the metaproteome of the cecal contents by mass spectrometry, which led to the identification of over 1000 host proteins and almost 400 proteins from the highly abundant *D. newyorkensis* (Supplemental Fig. 4C, Table 4). Comparing healthy and diseased mice, host proteins generally became more abundant in the cecal contents of the latter while microbial proteins followed the opposite trend (Supplemental Fig. 4C). As previously observed with the metabolomics data, this is consistent with the epithelial barrier maintaining separation between the host and the microbiota in the healthy state but then becoming compromised in the diseased state. The decreased abundance of microbial proteins in diseased mice likely indicates a decline in the relative levels or activities of microbes associated with healthy mice, such as *D. newyorkensis*.

We then searched for correlations between the composition of the microbiota and metabolite levels. In the cecal contents, one notable correlation involved 4-hydroxybutyrate (GHB), levels of which were negatively associated with *Erysipelotrichaceae* (OTU1) and positively associated with *Enterobacteriaceae* (OTU3) (Supplemental Fig. 4D). Although these correlations may indicate potential relationships between certain metabolites and microbes, disease severity also correlates with these variables, raising the possibility that other confounding factors, such as the host inflammatory response, could underlie the observed changes. Based on the analyses presented above, we show that inflammation disrupts the composition, activity, and compartmentalization of the gut microbiota and its associated molecules, all of which may affect the ability of the microbiota to supply carbon to the host for histone acetylation.

### Dietary fiber is a carbon source for hundreds of metabolites

Next, we compared mice treated with ^13^C-labeled versus unlabeled inulin to identify metabolite features containing carbon derived from dietary fiber. Across all tissues, we detected isotopic labeling on nearly 300 features (Fig. 4, Table 2). Molecules in the cecal contents were generally labeled to a greater extent than those in the cecal or colon tissue, consistent with a requirement for microbial processing before the labeled carbon becomes available to host cells. Only minor amounts of labeling surfaced in the more distally located liver tissue. As expected, ^13^C atoms from inulin appeared on intermediates from glycolysis and the TCA cycle, including pyruvate, alpha-ketoglutarate, succinate, and malate (Supplemental Fig. 5A, 5C). Consistent with previous literature, inulin underwent fermentation into SCFAs (Deroover *et al*, 2017; Butts *et al*, 2016) (Supplemental Fig. 5B). Roughly 12% of propionate demonstrated an isotopic shift to the m+2 isotopomer. Ethyl acetate, which overlaps in mass with butyrate, also displayed signs of ^13^C incorporation. Aside from its catabolic breakdown, inulin contributed to the biosynthesis of amino acids, nucleotide derivatives, and NAD (Supplemental Fig. 5D-F). In general, closely related molecules had similar isotopomer distributions, as highlighted by alpha-ketoglutarate, glutamate, and glutamine.

**Figure 4.**
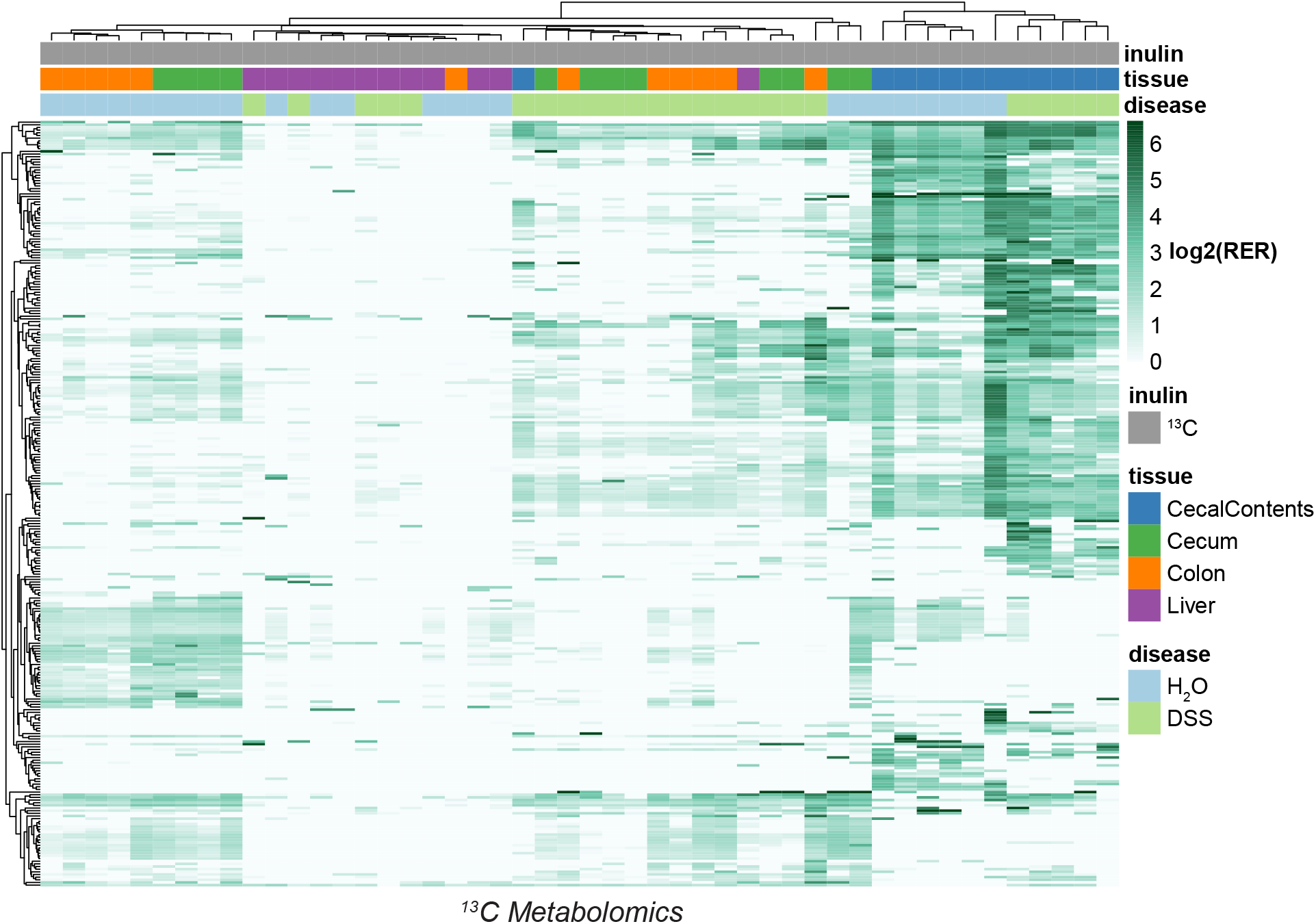
Dietary fiber is a carbon source for hundreds of metabolites. Heatmap of ^13^C isotope incorporation for 288 metabolite features classified as labeled. The color scale represents the log2 of the relative exchange rate (RER), except in cases where RER < 1, which were assigned a value of log2(1) = 0. NA values were replaced with the row minimum.

### Inflammation interferes with the contribution of fiber to host fatty acid metabolism

One of the more intriguing findings from the *in vivo* isotope tracing analysis involved molecules bearing long chain acyl groups, including free fatty acids, membrane lipids, and acylcarnitines. We noted the presence of inulin-derived carbon in several LCFAs, such as hydroxypentadecanoic acid, from the cecal contents of healthy mice (Fig. 5A). The isotopomer distributions for these LCFAs showed a pattern characteristic of the sequential transfer of two ^13^C atoms from fully labeled acetyl-CoA to an elongating acyl chain. LCFAs can be used anabolically for membrane biosynthesis or catabolically for fatty acid oxidation. Providing evidence for the former, we detected isotopic labeling of several lipids in both the cecal contents as well as the gut tissues (Fig. 5B). However, a portion of these acyl groups is likely transported into mitochondria for fatty acid oxidation since long-chain acylcarnitines from the cecum and colon also acquired the isotopic label (Fig. 5C). Interestingly, colitis curtailed the labeling of these acylated molecules (Fig. 5A-C). Overall, these results indicate that the microbiota utilizes dietary fiber to construct long-chain acyl groups and that inflammation interferes with either the transfer of these pre-assembled molecules into the cecal tissue or their *de novo* synthesis in gut epithelial cells using carbon flowing from the microbiota.

**Figure 5.**
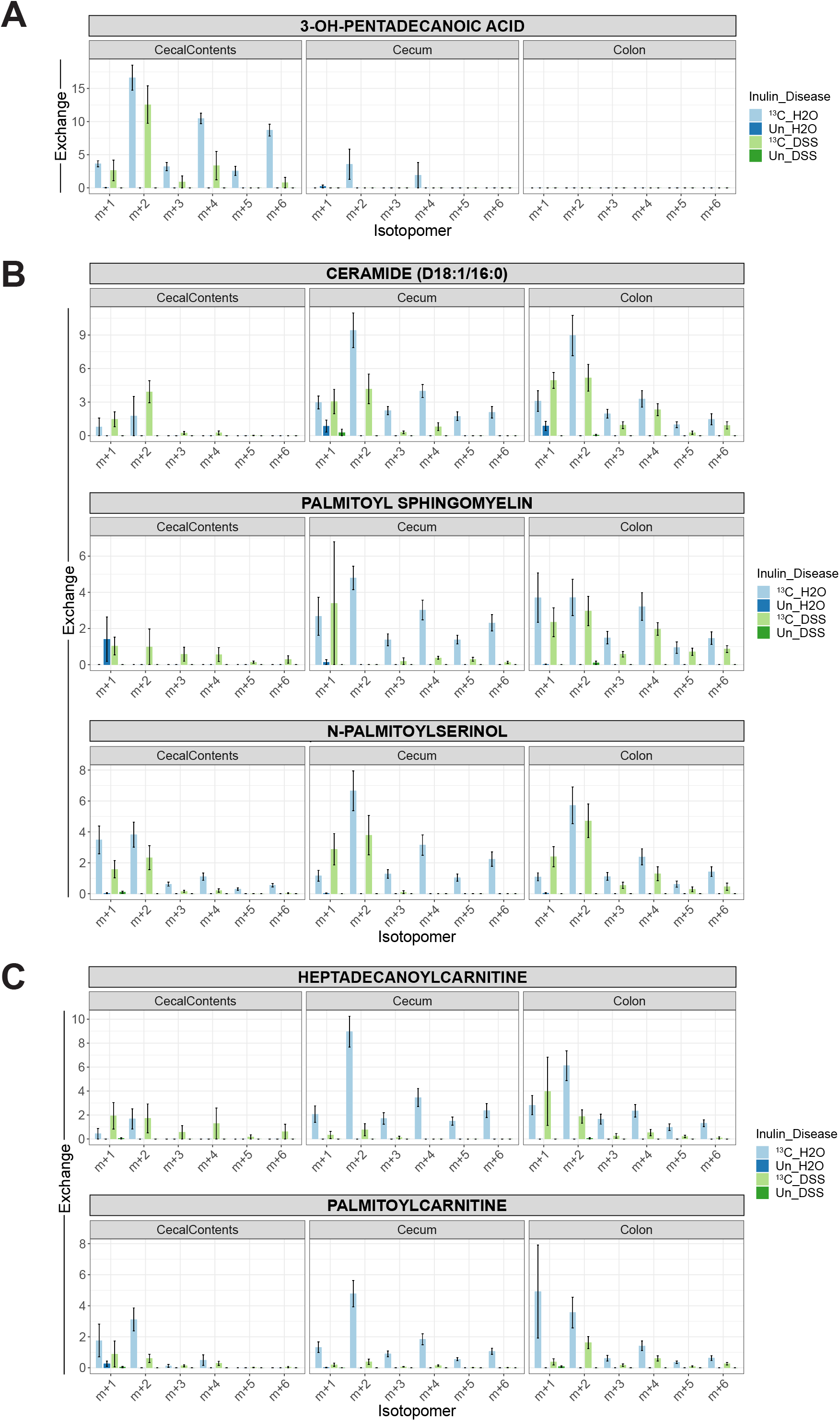
Inflammation interferes with the contribution of fiber to host fatty acid metabolism. A, B, C) Isotopomer distributions for metabolites with long-chain acyl groups, including LCFAs (A), membrane lipids (B), and acylcarnitines (C). Metabolites were isolated from the cecal contents, cecum, or colon. The monoisotopic peak (m+0) and isotopomers past the m+6 peak are omitted for clarity. The distributions are corrected for the relative abundances of naturally occurring isotopomers, meaning that unlabeled samples are expected to have no signal beyond the m+0 peak. Mean ± sem, n = 6.

### Inflammation suppresses cecal expression of genes supporting fatty acid metabolism

To correlate changes in our metabolomics data with changes in host gene expression, we performed RNA-seq in parallel. As shown in Fig. 6A, colitis had a greater impact on gene expression in the cecum, which displayed a transcriptional signature of inflammation based on upregulated expression of genes related to NLR and IL-23 signaling, such as *Il1b, Il6, Il17a, Il17f, Tnf*, and *Ccl2* (Supplemental Fig. 6A, Table 5). *S100a8* and *S100a9*, which together form calprotectin, a classical marker of colitis, were also upregulated. Concomitant with upregulated expression of inflammatory genes, diseased mice had downregulated expression of genes related to fatty acid metabolism (Fig. 6B), including members of the carnitine shuttle system (*Cpt1a*, *Cpt2*, *Crat*), acyl-CoA dehydrogenases (*Acads*, *Acadm*, *Acadvl*), and other enzymes directly responsible for catabolizing fatty acids into acetyl-CoA (*Ehhadh*, *Hadh*, *Hadha*, *Hadhb*). Colitis also reduced the expression of peroxisomal genes (Supplemental Fig. 6B). Like mitochondria, peroxisomes are capable of fatty acid oxidation and have particular importance in the catabolism of certain fatty acids, such as VLCFAs, branched chain FAs, and dicarboxylic FAs. These changes in gene expression may reflect changes in the relative proportions of epithelial cells and inflammatory cells in the cecum. Overall, we find that colitis disrupted the expression of genes supporting cecal fatty acid metabolism, correlating with the reduced flux of inulin-derived carbon to acylated molecules.

**Figure 6.**
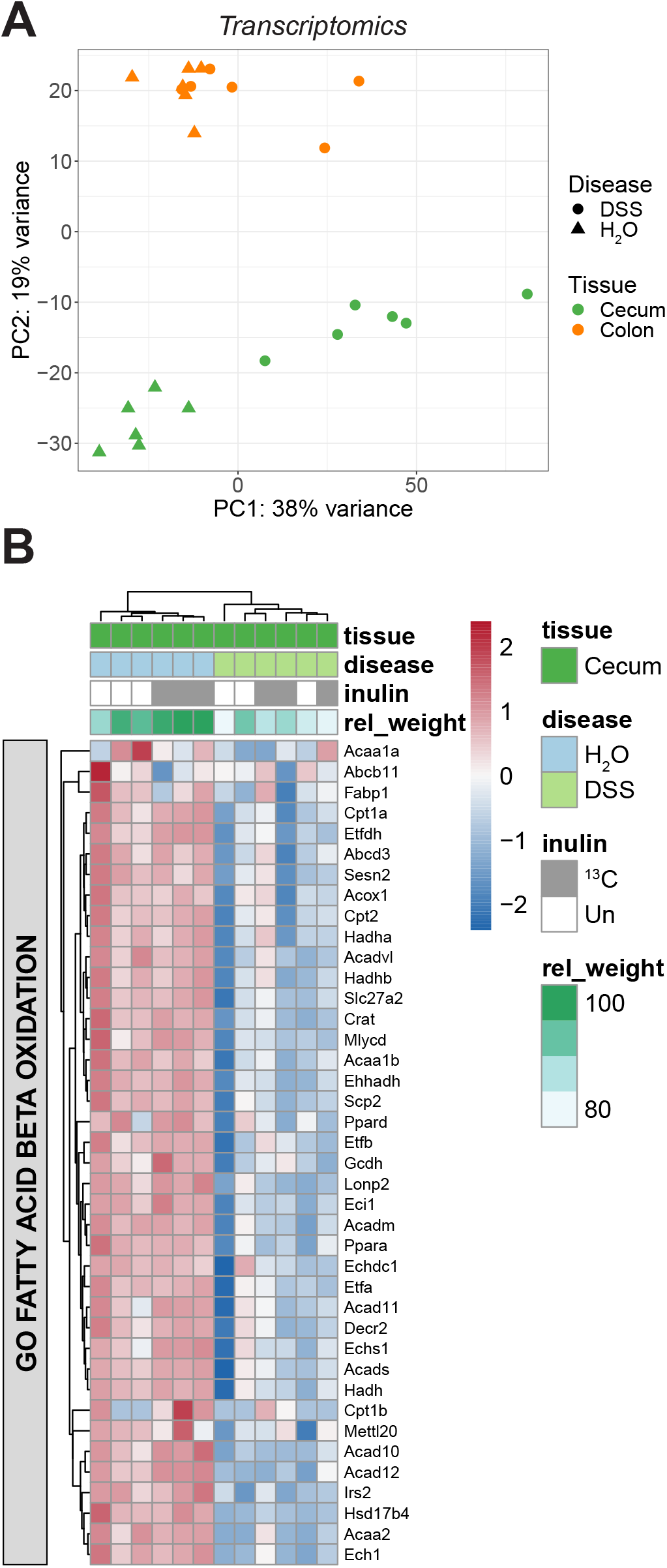
Inflammation suppresses cecal expression of genes supporting fatty acid metabolism. A) PCA of whole cecum and colon tissue based on 15,825 transcriptomic features (n = 6, 3 each from Un and ^13^C inulin groups). B) Heatmap of core enrichment genes related to fatty acid oxidation based on GSEA of the cecal transcriptome. The relative weight change for each mouse is also indicated at the top.

## DISCUSSION

Host-microbiota interactions are known to influence many physiological processes in the gut. Here, we initially focused on the impact of the microbiota on host epigenetics in the cecum and colon, finding that the lack of this important community leads to a reduction in histone H4 acetylation across the genome except at transcription start sites. Through isotope tracing experiments with butyrate in cell culture and dietary fiber in mice, we demonstrate that butyrate and other products of microbial metabolism act as carbon sources for histone acetylation. Using a metabolomics approach, we also show a broader contribution of microbial metabolism to host physiology, as evident by our tracking of the isotope label to acylated molecules, including acylcarnitines and membrane lipids. We also document, to our knowledge, the first direct evidence that this microbiota-dependent metabolic link to histone acetylation operates *in vivo*. Finally, we observe that inflammation interferes with carbon flux from the microbiota to host fatty acid metabolism.

Many studies have demonstrated that the microbiota promotes histone acetylation (Furusawa *et al*, 2013; Krautkramer *et al*, 2016). Our results showing higher levels of acetylation on histone H4 in conventional versus germ-free mice agree with those of a previous publication that involved a similar MS-based analysis (Krautkramer *et al*, 2016). The positive effect of the microbiota on histone acetylation has been linked to the fermentation of dietary fiber, which produces SCFAs like butyrate (Krautkramer *et al*, 2016; Furusawa *et al*, 2013). However, butyrate may not accumulate to significant levels in gut epithelial cells since they are adapted to derive their energy from SCFAs through fatty acid oxidation. Thus, rather than supporting histone acetylation via inhibition of a negative regulator (HDACs), the microbiota may activate a positive regulator (HATs) via provision of substrates that feed the production of acetyl-CoA, the donor substrate for HATs. These two mechanisms of butyrate-mediated promotion of histone acetylation were demonstrated by experiments showing that silencing of ATP citrate lyase (ACLY) prevents increases in histone H3 acetylation at low (<0.5 mM) but not high (>2 mM) doses of butyrate (Donohoe *et al*, 2012). Earlier studies in cell culture suggested that butyrate may act as a carbon donor for histone acetylation based on incorporation of ^14^C into histones and the detection of ^13^C-labeled acetate hydrolyzed from histones (Donohoe *et al*, 2012; Andriamihaja *et al*, 2009). Our isotope tracing analysis with ^13^C-labeled butyrate in cell culture, combined with the high-resolution analysis of histone peptides by mass spectrometry, extends these results and shows unequivocally that the acetyl groups of histones contain carbon derived from butyrate. Importantly, using ^13^C-labeled fiber in a more physiologically relevant model, we obtained direct evidence that this microbiota-dependent metabolic link to histone acetylation operates *in vivo* as well. The extent to which butyrate functions as an inhibitor of HDAC activity versus a supplier of acetyl-CoA in support of HAT activity remains unclear but likely relates to its concentration and the rate at which cells metabolize it (Donohoe *et al*, 2012). Given the recent and unexpected observation that HDAC activity is higher in intestinal epithelial cells from conventional, butyrate-sufficient mice versus germ-free, butyrate-deficient mice, HDAC inhibition may not be the primary activity of butyrate in the gut (Wu *et al*, 2020).

Although the global decrease in H4 acetylation in germ-free mice that we observe has been noted previously (Krautkramer *et al*, 2016), we have expanded upon these earlier findings by performing ChIP-seq and RNA-seq to ascertain where this reduction occurs in the genome and the consequences for gene expression. These results showed a loss of H4 acetylation distributed across the genome, particularly at gene bodies. This finding mirrors what has been recently described for H4 acetylation in cells treated with an HDAC inhibitor, causing an overall elevation in H4 acetylation with some of the most significant increases occurring in gene bodies (Slaughter *et al*, 2021). Despite the global loss of H4 acetylation in germ-free mice, we find that this histone modification becomes more pronounced at transcription start sites. However, these epigenetic changes appear to have only minor effects on transcription since we find relatively small and balanced numbers of upregulated and downregulated genes in germ-free mice. This is consistent with a prior analysis comparing chromatin accessibility in conventional and germ-free mice, which found that the anatomical location along the intestinal tract but not colonization status influenced the epigenetic landscape (Camp *et al*, 2014). Instead, variations in gene expression due to colonization status were proposed to arise from microbiota-dependent transcription factors acting on a pre-configured epigenetic landscape established during early development. While some differences in the genomic patterns of H3K4me1 and H3K27ac were noted in a subsequent study that compared epithelial cells from the jejunum of germ-free and formerly germ-free mice, a more remarkable observation was the increased binding of the nuclear receptors HNF4A and HNF4G to areas of chromatin in germ-free mice that had similar patterns of accessibility and histone modifications, again emphasizing an important contribution of stimulus-dependent transcription factors in controlling gene expression (Davison *et al*, 2017). In this regard, it is especially noteworthy that microbial metabolites can act as ligands for nuclear receptors (S. Ranhotra *et al*, 2016; Klepsch *et al*, 2019; Duszka & Wahli, 2018).

The contribution of the microbiota to histone acetylation through metabolism, as revealed by our isotope tracing studies, prompted us to examine isotope incorporation more broadly using metabolomics to determine how microbial metabolism of dietary fiber supplies precursors for host cell metabolism. The idea that the microbiota supports host metabolism in the gut has been suggested by earlier studies describing the transfer of isotope-labeled carbon from butyrate to CO_2_ in cultures of primary colonocytes as well as established cell lines (Roediger, 1980b, 1980a; Ahmad *et al*, 2000; Chapman *et al*, 1994; Andriamihaja *et al*, 2009; Leschelle *et al*, 2000; Donohoe *et al*, 2011). Some studies additionally tracked the isotope label to acetyl-CoA, histones, and lipids (Leschelle *et al*, 2000; Andriamihaja *et al*, 2009; Kaiko *et al*, 2016). Consistent with the microbiota supporting host metabolism, germ-free mice display signs of energy deprivation in the colon as indicated by lower levels of ATP and a decreased NADH/NAD+ ratio, likely resulting from less flux through the TCA cycle and electron transport chain (Donohoe *et al*, 2011). Notably, this energy deficit could be rescued by introducing a normal microbiota or a single butyrate-producing organism into germ-free mice.

Bringing clinical significance to this metabolic connection, similar observations of impaired butyrate oxidation and mitochondrial function have been reported in the case of ulcerative colitis in humans and DSS-induced colitis in mice (Roediger, 1980b; Ahmad *et al*, 2000; Chapman *et al*, 1994; Smith *et al*, 2021; Haberman *et al*, 2019; De Preter *et al*, 2012). Building on these *in vitro* and *ex vivo* analyses, we conducted isotope tracing *in vivo* with ^13^C-labeled fiber and obtained definitive evidence of carbon transfer from the microbiota to the host under physiological conditions. A recent study similarly employed ^13^C-labeled fiber to interrogate the metabolic pathways of the microbiota using an *in vitro* system (Deng *et al*, 2021), which affords more control over the labeling conditions but cannot address potential interactions with host tissues like our approach. Among our most significant findings, we detected incorporation of the isotope label into the acyl chains of palmitoylcarnitine and membrane lipids. These results support the notion that the microbiota acts as a carbon source for acetyl-CoA, which can then be used by gut epithelial cells to drive the TCA cycle, acetylate proteins, or construct long-chain acyl-CoA for lipid synthesis (Fig. 7). Our observation that DSS-induced colitis interfered with the isotopic labeling of these acylated molecules aligns with earlier work correlating colitis with deficient fatty acid metabolism. A major question is whether this deficiency is a primary factor in the development of ulcerative colitis. Insufficient energy production could compromise the function of epithelial cells, causing a breakdown of the epithelium and a consequent immune response to microbial products that leak into host tissues. Although it is unclear what might trigger an initial decline in fatty acid metabolism, antibiotic usage, low-fiber diets, or infections could all reduce the ability of the microbiota to produce the SCFAs that fuel epithelial cell metabolism (Litvak *et al*, 2018). Excessive regeneration of the epithelium may also favor the non-oxidative metabolism of undifferentiated cells (Litvak *et al*, 2018; Lopez *et al*, 2016). On the other hand, deficient metabolism could represent a promoting factor or even a simple consequence rather than a primary cause of ulcerative colitis since the microbial communities associated with the disease and an inflamed epithelium may have respectively lower capacities for SCFA production and fatty acid oxidation.

**Figure 7.**
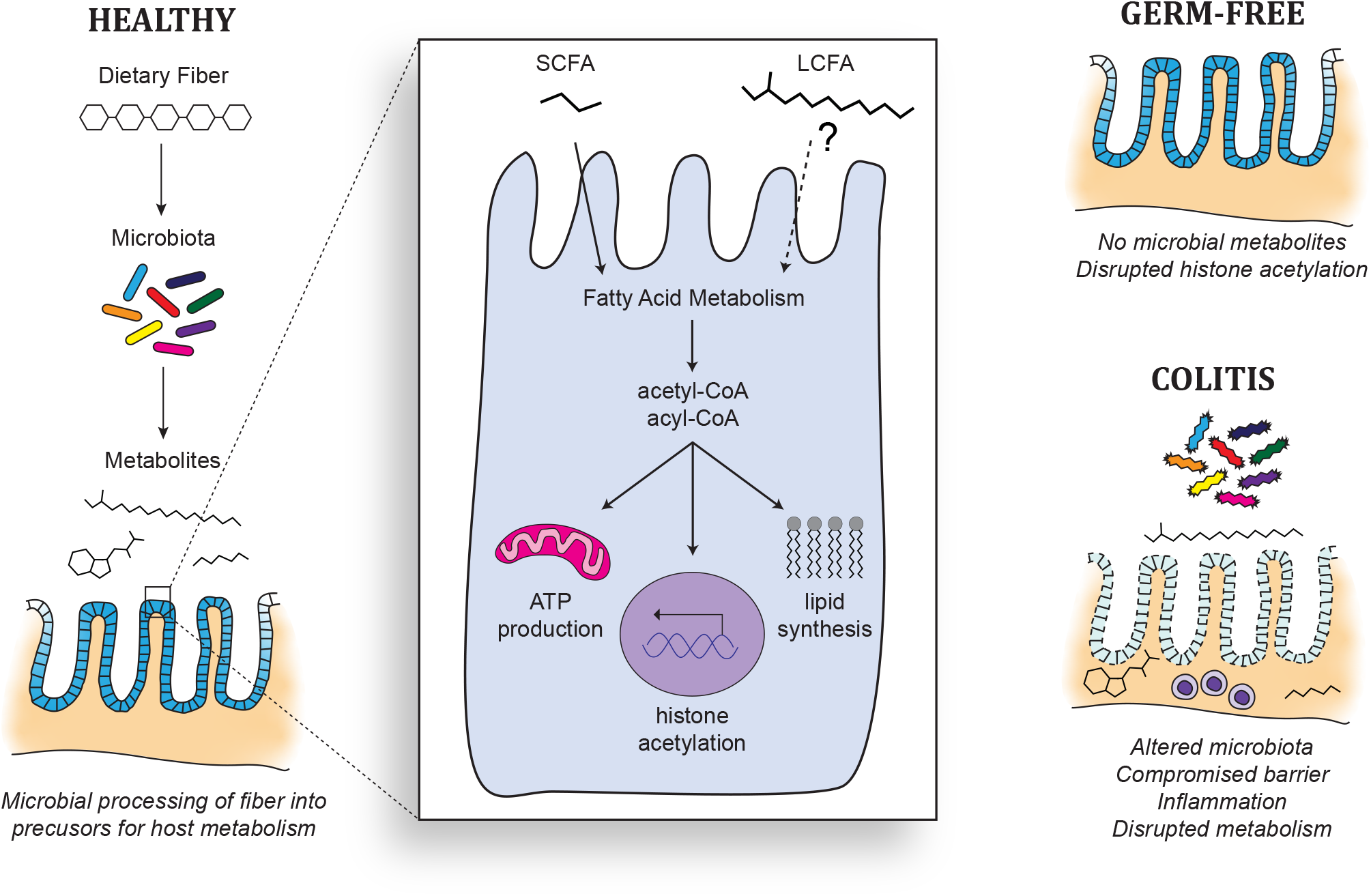
Proposed model linking dietary fiber and the microbiota to fatty acid metabolism in gut epithelial cells. Dietary fiber is processed by the microbiota to generate precursors, such as short-chain fatty acids, for fatty acid metabolism in host epithelial cells. The resulting acetyl- and acyl-CoA can be utilized to acetylate histones, synthesize lipids, or fuel the TCA cycle for ATP production. Long-chain fatty acids from the microbiota may also contribute to these processes. In the absence of a microbiota, GF mice must rely on other sources of metabolic precursors. Similarly, inflammation interferes with carbon flow from fiber to the host via the microbiota.

The labeling of long-chain acyl groups also raises the question of their origin and whether LCFAs might complement SCFAs as energy sources in the gut. One possibility accounting for these labeled acyl groups involves the oxidation of labeled SCFAs to acetyl-CoA, which could then be used by host cells to synthesize LCFAs. Alternatively, the labeled acyl groups may come directly from microbial lipids or LCFAs, which could be made available to host cells through membrane vesicles or the action of secreted phospholipases. In support of LCFAs originating directly from the microbiota, a recent study imaged the transfer of lipids from bacteria to mammalian cells (Johnson *et al*, 2020). Besides their assimilation into host membranes, these microbial lipids could also be oxidized for energy, as indicated by our detection of labeled palmitoylcarnitine. Indeed, the same energy deficiency observed in germ-free mice (Donohoe *et al*, 2011) has not been described, to our knowledge, in mice lacking *Scad* activity, which is necessary for the utilization of SCFAs. This suggests that SCFAs may not constitute the only pool of microbiota-dependent carbon sources in the gut.

Sequencing approaches have facilitated our understanding of the community structure and coding potential of the microbiota, which is estimated to exceed the gene content of the human genome by a factor of 150x (Qin *et al*, 2010). Further insight into how the microbiota influences host physiology necessitates the complementary study of microbial proteins and small molecules, representing the functional output of the microbiota. Overall, the current study contributes to our knowledge of host-microbiota interactions by using isotopic labeling to demonstrate a connection between microbial fermentation of dietary fiber and host fatty acid metabolism, which may be important for gut homeostasis.

## MATERIALS AND METHODS

### Animal studies

All experiments were performed with approval from the Institutional Animal Care and Use Committee. Unless otherwise noted, mice were C57BL/6J females from Jackson Labs, age 7-8 weeks. Germ-free mice were provided by the Penn Gnotobiotic Mouse Facility. Animals were euthanized by CO_2_ asphyxiation followed by cervical dislocation. Colons and ceca were dissected, flushed with sterile phosphate-buffered saline (PBS), splayed open longitudinally, and washed again with PBS for use in epithelial cell isolation. For the inulin labeling timecourse, mice were acclimated to an irradiated AIN76a diet (Research Diets, D10001) for 3 days and then orally gavaged with 100 mg of U-^13^C-labeled inulin (Isolife), dissolved in 100 μl PBS, once per day for up to 3 days. Control mice received PBS alone or unlabeled inulin. One day after the last gavage, colon tissue was harvested for epithelial cell isolation. For the antibiotic experiment, mice were acclimated to Splenda water (10 mg/ml) over 3 days and then treated with or without antibiotics (1 mg/ml ampicillin, 1 mg/ml neomycin, 0.5 mg/ml metronidazole, 0.5 mg/ml vancomycin) in the Splenda drinking water and switched to the AIN76a diet. On days 5 and 6, mice received an oral gavage of 10, 50, or 100 mg of U-^13^C-inulin or 100 mg unlabeled inulin. On day 7, colon tissue was harvested for epithelial cell isolation. For the colitis experiment, mice (8-10 wks) were acclimated to the AIN76a diet over the preceding 3 days and then had colitis induced with 2% dextran sodium sulfate (MP Bio, 36-50 kDa) in the drinking water on day 0. On days 6 and 7, mice received an oral gavage of 100 mg of U-^13^C-labeled or unlabeled inulin (IsoLife) dissolved in 200 μl PBS. On day 8, mice were sacrificed by cervical dislocation after a 4 hour fast. Livers, colons, ceca, and cecal contents were harvested and flash frozen in liquid nitrogen. An aliquot of the cecal contents was submitted to the Weill Cornell Microbiome Sequencing Core, which performed 16S sequencing and analysis.

### Epithelial cell isolation

To prepare epithelial cell pellets, colons and ceca were incubated in PBS containing 30 mM EDTA and 1.5 mM DTT for 20 minutes at 4°C with occasional inversion. Tissue was subsequently transferred to PBS containing 30 mM EDTA, incubated for 10 mins at 37°C, and shaken vigorously for 1-2 minutes until the mesenchyme separated from the epithelium. The mesenchyme was removed and epithelial cells were pelleted at 1500 rpm for 5 minutes at 4°C and washed in PBS. In some cases, cell pellets were further incubated with dispase (Corning, 0.3 U/ml) in PBS (100 μl dispase per 50 ml PBS) at 37°C for 10 mins and then quenched with 1/10 volume of FBS containing DNase (10 ul DNase per ml FBS) followed by washing in PBS. Cell pellets were flash frozen in liquid nitrogen and stored at −80°C until use. For ChIP-seq analysis, cells were cross-linked prior to freezing as indicated below.

### Cell culture

Standard conditions for the culture of HEK293 and Caco2 cells consisted of DMEM supplemented with 10% FBS or FetalPlex and 1X penicillin/streptomycin in a humidified incubator at 37°C. Starved conditions consisted of DMEM without glucose or pyruvate (but with glutamine) supplemented with 1% dialyzed FBS and antibiotics. For isotope labeling experiments, cells were seeded in 5 or 10 cm dishes, allowed to grow for 24-48 hours, and then treated with 1 mM unlabeled or U-^13^C-labeled sodium butyrate (Sigma). Cell monolayers were rinsed with PBS, trypsinized, washed again in PBS, and flash frozen in liquid nitrogen.

### Histone extraction and analysis by LC-MS/MS

Histones were isolated from the nuclei of frozen cell pellets by acid extraction and derivatized with propionic anhydride similarly as described previously (Lund *et al*, 2019). Briefly, cell pellets were resuspended in nuclear isolation buffer (NIB) containing 15 mM Tris pH 7.5, 15 mM NaCl, 60 mM KCl, 5 mM MgCl_2_, 1 mM CaCl_2_, and 250 mM sucrose supplemented with 1 mM DTT, 10 mM sodium butyrate, 500 μM AEBSF, 5 nM microcystin, and 0.2% NP-40 Alternative. After allowing lysis to proceed on ice for 10 mins, nuclei were pelleted at 500 x g for 5 mins at 4°C, washed twice in NIB without detergent, and then extracted in 0.4 N H_2_SO_4_ for 2-4 hrs at 4°C with rotation. Insoluble debris was pelleted by centrifugation at 3400 x g for 5 mins at 4°C and proteins were precipitated overnight on ice by adding 1 volume of trichloroacetic acid to 3 volumes of supernatant. Precipitated proteins were pelleted as above, washed in 0.1% HCl in acetone and then acetone, and allowed to air dry before resuspending in 100 mM NH_4_HCO_3_. For derivatization, 1 volume of 25% propionic anhydride in 2-propanol was added to 2 volumes of sample containing 5-20 μg of protein. Ammonium bicarbonate salt was also added for buffering purposes. After incubation for 15 mins at 37°C, samples were dried in a speed vac and derivatized a second time as above. Derivatized samples were then resuspended in 100 mM NH_4_HCO_3_ and digested overnight at room temperature with 1 μg trypsin per 20 μg of protein. After two additional rounds of derivatization, digested peptides were desalted with C18 stage tips for analysis by mass spectrometry.

Histone peptides were resolved with EasyLC 1000 or Dionex UltiMate 3000 LC systems fitted with 75 μm i.d. x 15-20 cm fused silica columns (Polymicro Tech) packed with ReproSil-Pur 120 C18-AQ (3 μm, Dr. Maisch GmbH) and connected in line with a mass spectrometer (Thermo Elite, Velos, Fusion, QE, or QE-HF). The chromatography conditions generally consisted of a linear gradient from 5 to 33% solvent B (0.1% formic acid in 80% acetonitrile) in solvent A (0.1% formic acid in water) over 45 mins and then 33 to 98% solvent B over 5 mins at a flow rate of 300 nl/min. The mass spectrometer was programmed for data-independent acquisition (DIA). One acquisition cycle consisted of a full MS scan, 8 DIA MS/MS scans of 50 m/z isolation width starting from 325 m/z, a second full MS scan, and 8 more DIA MS/MS scans to reach 1125 m/z. Typically, full MS scans were acquired in the Orbitrap mass analyzer across 300-1100 m/z at a resolution of 60,000 in positive profile mode with a maximum injection time of 100 ms and an AGC target of 2e5. MS/MS data from CID or HCD fragmentation was collected in the ion trap (when available) or the Orbitrap. These scans typically used an NCE of 30, an AGC target of 1e4, and a maximum injection time of 50 ms. Spectral data, including isotope incorporation, were analyzed with EpiProfile (Yuan *et al*, 2015) and further processed in R to filter for peaks with consistent retention times.

### Metabolomics

Frozen tissues were lyophilized and then homogenized in 80% methanol with bead beating (Precellys homogenizer, Bertin Technologies) at a target concentration of 5-10 mg tissue per 500 μl of solvent. For tissues with limited quantities, a minimum of 250 μl solvent was used. For lyophilized tissues exceeding 10 mg, bead beating was used to ground them into a dry powder, of which roughly 10 mg was used for homogenization. Aliquots of the homogenates were treated with additional methanol and then centrifuged to precipitate proteins. The supernatants were dried under nitrogen and then resuspended in HPLC mobile phases for resolution on C18 or HILIC columns in line with a Thermo Orbitrap ID-X mass spectrometer. For C18 chromatography, extracted samples were injected onto a Waters Acquity BEH C18 column (2.1 x 150 mm, 1.7 μm) at 65°C using a 13 min linear gradient from 99% solvent A (0.1% formic acid in water) to 100% solvent B (acetonitrile with 0.1% formic acid) at a flow rate of 0.6 ml/min on a Thermo Vanquish UHPLC. The mass spectrometer was run in ESI+/- modes from 60-1000 Da at a resolution of 120,000. For HILIC, extracted samples were injected onto an Agilent Poroshell HILIC-Z column (2.1 x 100 mm, 1.9 μm) at 30°C using a 10 min linear gradient from 95% solvent B (acetonitrile) to 70% solvent A (water with 10 mM ammonium acetate pH 9 and 0.1% medronic acid) at a flow rate of 0.4 ml/min to separate highly polar metabolites.

CompoundDiscoverer (CD) was used for data processing and isotope incorporation analysis. Additional analyses were performed in R. For the analysis of overall metabolite levels, peak areas from CD were first multiplied by a correction factor to standardize extraction conditions to 20 mg/ml and then normalized by log2 transformation. The relative exchange rate (RER) calculated by CD was used to identify features with ^13^C incorporation based on consistently higher RER in labeled versus unlabeled conditions and a low background RER in the unlabeled conditions (log2[labeled/unlabeled] >1; p < 0.01 from unpaired, two-tailed t-test; maximum mean RER for unlabeled < 1; maximum median RER for labeled >2). In cases where the log2 FC or p-value was undefined, a feature was considered as labeled if the mean RER for unlabeled conditions within a given tissue was zero and the mean RER for the labeled condition was greater than 2.

### ChIP-seq and RNA-seq

Cross-linking was performed by treatment with 1% formaldehyde in PBS for 5 mins at room temperature, followed by quenching with 125 mM glycine for an additional 5 mins at room temperature. Cross-linked pellets were then washed in PBS and flash frozen until use. Nuclear lysates were prepared by resuspending cells in lysis buffer 1 (50 mM HEPES-KOH pH 7.5, 140 mM NaCl, 1 mM EDTA, 10% glycerol, 0.5% NP-40, 0.25% Triton X-100, 1x protease inhibitors) and rotating for 10 mins at 4°C. After centrifugation at 3000 rpm for 5 mins at 4°C, the cell pellet was resuspended in lysis buffer 2 (10 mM Tris-HCl pH 8, 200 mM NaCl, 1 mM EDTA, 0.5 mM EGTA, 1x protease inhibitors) and rotated for 10 mins at room temperature. Cells were pelleted and then resuspended in lysis buffer 3 (10 mM Tris-HCl pH 8, 100 mM NaCl, 1 mm EDTA, 0.5 mM EGTA, 0.1% sodium deoxycholate, 0.1% N-lauroylsarcosine, 1x protease inhibitors) for chromatin shearing with a Covaris S220 focused ultrasonicator (8.4W, 200 peak power, 200 cycles/burst, 900 seconds) in a 4°C water bath. After sonication, Triton X-100 was added to 1% final concentration and the lysate was cleared by centrifugation at top speed in a microfuge for 10 mins at 4°C. Magnetic protein G beads were washed with blocking solution (0.5% BSA in PBS) and then incubated with rabbit antiserum against acetylated histone H4 (Millipore 06-866) or normal rabbit anti-serum in blocking solution for several hours at 4°C with rotation. Beads were then washed in blocking solution and then added to nuclear lysates containing a spike-in of *Drosophila* S2 chromatin (cells generously provided by P. Pascual-Garcia), prepared as above. After overnight incubation at 4°C, beads with captured chromatin were washed with 3 x 1 ml RIPA wash buffer (50 mM HEPES-KOH pH 7.5, 500 mM LiCl, 1 mM EDTA, 1% NP-40, 0.7% sodium deoxycholate) and 1 x 1 ml of final wash buffer (10 mM Tris-HCl pH 8, 50 mM NaCl, 1 mM EDTA). Beads were eluted with 200 μl elution buffer (50 mM Tris-HCl pH 8, 10 mM EDTA, 1% SDS) for 30 mins at 65°C. The supernatant was then incubated at 65°C overnight to reverse cross-links. After dilution with one volume of TE, RNase was added to 0.2 mg/ml and incubated at 37°C for 2 hrs. Proteinase K was then added to 0.2 mg/ml and incubated at 55°C for 2 hrs. DNA was purified by phenol-chloroform extraction with subsequent ethanol precipitation. Purified DNA was resuspended in 10 mM Tris pH 8 and prepared for sequencing using the NEBNext Ultra II DNA library kit for Illumina (New England Biolabs). Paired-end sequencing with 75 cycles and a 6-cycle index read was performed on the Illumina NextSeq500 system.

Total RNA was extracted from epithelial cell pellets and frozen tissues using the RNeasy kit (Qiagen). The QIAshredder column was used for epithelial cell pellets. For frozen tissues from the colitis study, homogenization was carried out in the provided lysis buffer using bead beating (Precellys homogenizer, Bertin Technologies). The NEBNext Ultra or Ultra II Directional RNA Library Prep Kit for Illumina with the mRNA magnetic isolation module (New England Biolabs) was used to prepare sequencing libraries from 1 μg of total RNA.

Sequencing reads were aligned with STAR (v2.5.2a) (Dobin *et al*, 2013) (ChIP-seq parameters: --outFilterMultimapNmax 20 --outFilterMismatchNmax 999 --alignMatesGapMax 1000000; RNA-seq parameters: --outFilterType BySJout --outFilterMultimapNmax 20 --alignSJoverhangMin 8 -- alignSJDBoverhangMin 1 --outFilterMismatchNmax 999 --alignIntronMax 1000000) to the mm10 genome assembly or the mm10 genome concatenated with the *Drosophila* dm6 genome in the case of the ChIP with the exogenous spike-in. Alignments were filtered for uniquely mapping reads. Transcriptome data was analyzed quantitatively with HTSeq (Pyl *et al*, 2014) and DESEQ2 (Anders & Huber, 2010). Pathway analysis was performed using GSEA (Subramanian *et al*, 2005) or the ClusterProfiler and ReactomePA packages in R (Yu *et al*, 2012; Yu & He, 2016). For ChIP, normalization to the exogenous reference chromatin in lieu of sequencing depth was performed as described previously (Fursova *et al*, 2019). Briefly, downsampling factors, corrected for any variations in reference chromatin abundance in the input samples, were calculated and applied to each sample such that equal numbers of *Drosophila* reads were obtained across all samples with the largest factor being equal to 1. BigWig files were generated using an in-house script. DeepTools was used for additional analysis and visualization.

### Metaproteomics

Protein precipitates from the metabolite extractions of cecal contents were resolubilized in 8 M urea, 0.1 M NaCl, and 50 mM Tris pH 8 supplemented with protease inhibitors. Insoluble debris was pelleted by centrifugation. Roughly 20 μg of soluble protein was reduced with 5 mM DTT for 30 mins at room temperature and then alkylated with 20 mM iodoacetamide for 30 mins at room temperature in the dark. After adding four volumes of 0.1 M ammonium bicarbonate, proteins were digested with 1 μg trypsin overnight at 37°C and then desalted with C18 stage tips for mass spectrometry analysis. As for the histone analysis, peptides were separated by reverse-phase chromatography with an online EasyLC1000 nano-LC system. The gradient consisted of 2 to 30% solvent B (80% acetonitrile with 0.1% formic acid) over 52 mins, 30 to 60%solvent B over 24 mins, and 60 to 90% solvent B over 2 mins at a flow rate of 300 nl/min. Water with 0.1% formic acid served as solvent A. The mass spectrometer (Thermo QE) was operated in data-dependent acquisition (DDA) mode. A full scan in positive profile mode was acquired over the range of 300-1400 m/z at a resolution of 70,000, AGC target of 1e6, and maximum IT of 100 ms. The top 15 precursor ions were selected for HCD fragmentation at NCE 30 and MS/MS scans were collected at a resolution of 17,500 with an AGC target of 1e5, a maximum IT of 50 ms, and an isolation width of 2.0 m/z in centroid mode. Only ions with charge states between +2 and +6 were considered. Dynamic exclusion was set to 40 seconds. The minimum AGC target was set to 1e3. Data was analyzed with ProteomeDiscoverer using a custom metaproteome constructed by concatenating the mouse proteome with the proteomes of bacteria from the most abundant OTUs based on 16S sequencing. For database searches, trypsin was set as the protease with up to 2 missed cleavages. The mass tolerances were 10 ppm for precursors and 0.02 Da for fragments. Carbamidomethylation (+57.021 Da to Cys) was specified as a fixed modification while oxidation (+15.995 Da to Met), carbamylation (+43.006 Da to Lys and peptide N-termini), acetylation (+42.011 Da to protein N-termini), methionine loss (−131.040 Da to protein N-termini), and methionine loss with acetylation (−89.030 Da to protein N-termini) were set as variable modifications. The Percolator node was used to control for false-positive PSMs. The data was further analyzed in R with the MSStats package (MacLean *et al*, 2014).

## ACKNOWLEDGEMENTS

Funding is generously acknowledged from the Crohn’s and Colitis Foundation (RFA 598467 to P.J.L), the Crohn’s and Colitis Foundation Microbiome Initiative, the PennCHOP Microbiome Program, The Rockefeller University (C.D.A.), the Shapiro-Silverberg Fund for the Advancement of Translational Research at The Rockefeller University (L.A.G.), the St. Jude Children’s Research Hospital Chromatin Consortium, the Host-Microbial Analytic and Repository Core of the Center for Molecular Studies in Digestive and Liver Diseases (P30 DK050306), and the National Institutes of Health (T32CA009140 to P.J.L, F32GM134560 to L.A.G., P01CA196539 and R01AI118891 to B.A.G., R01GM40922 to C.D.A.). The authors wish to thank Johayra Simithy, Sophie Trefely, Kevin Janssen, Simone Sidoli, Greg Donahue, Pau Pascual-Garcia, Nathaniel W. Snyder, and Meenakshi Bewtra for assistance. P.J.L. graciously acknowledges Warren Pear for additional support and mentoring.

## CONTRIBUTIONS

P.J.L. participated in project conception, performed experiments, analyzed data, and drafted the original manuscript. L.G., M.L., S.A.S, L.C., E.S.F., Y.S., M.L., M.S.K., and C.P. performed experiments and assisted with sample preparation. C.D.A., G.D.W, and B.A.G. participated in project conception and supervision. All authors assisted with review and editing of the manuscript.

## SUPPLEMENTAL FIGURE LEGENDS AND TABLES

**Supplemental Figure 1.**
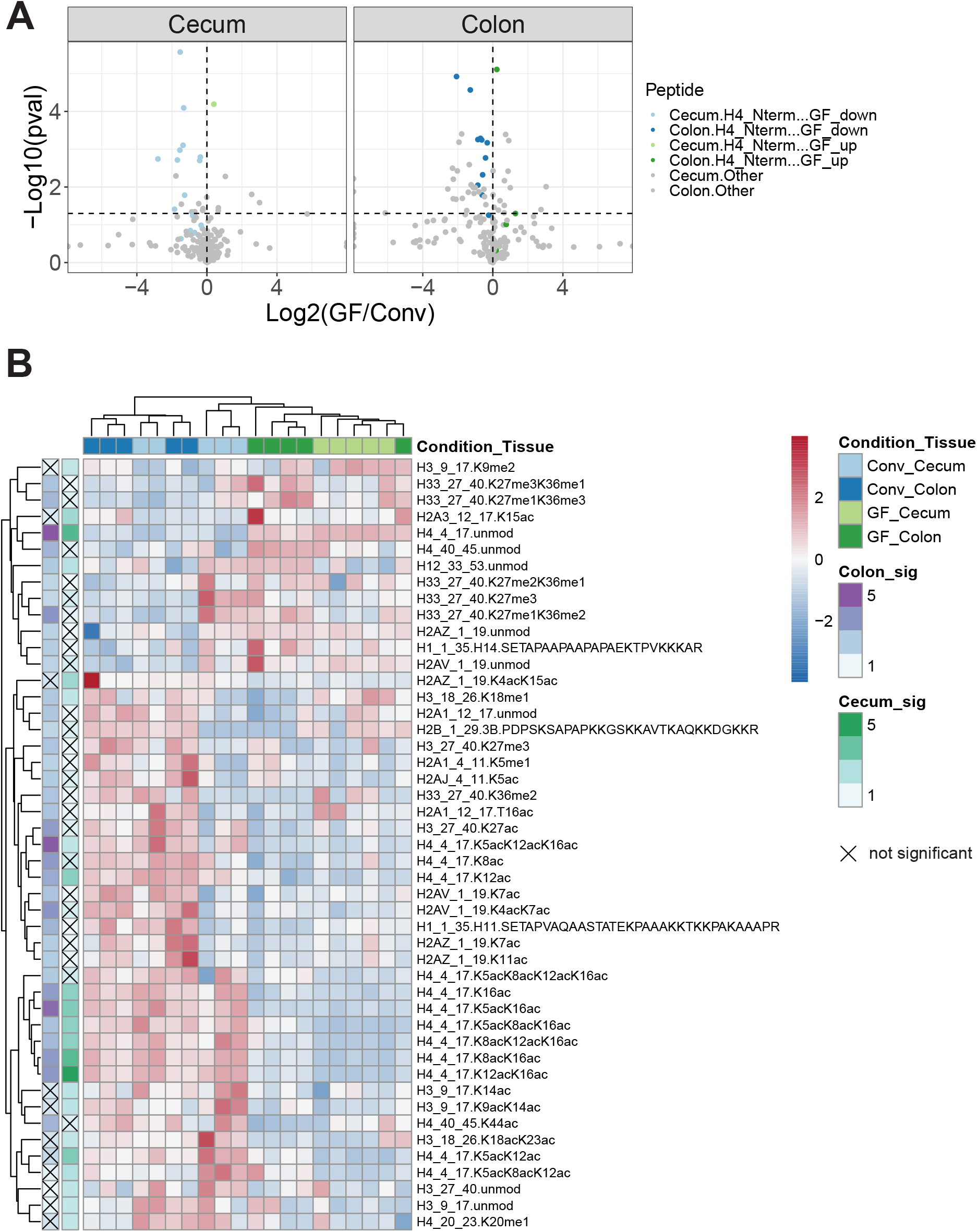
Global profiling of histone modification patterns in the cecum and colon of germ-free versus conventional mice. A) Volcano plot of fold-change versus statistical significance for uniquely modified histone peptides across each tissue (cecum: 198, colon: 199). Peptides from the N-terminal tail of histone H4 (4-17 aa) are shown in green and blue while all other peptides are shown in gray. A horizontal dashed line is shown at -log10(0.05). An unpaired, two-tailed t-test without the assumption of equal variances was used for statistical significance (n = 5). B) Heatmap of histone peptides with significant changes (p < 0.05 by unpaired, two-tailed t-test) in their relative abundance in the ceca or colons of GF versus Conv mice. Each peptide is annotated with its parent histone protein, its starting and ending amino acid positions, and its post-translational modifications (unmod: unmodified, ac: acetyl, me1: mono-methyl, me2: dimethyl, me3: tri-methyl). The -log10(pval) in each tissue (Colon_sig, Cecum_sig) is represented in boxes on the left side.

**Supplemental Figure 2.**
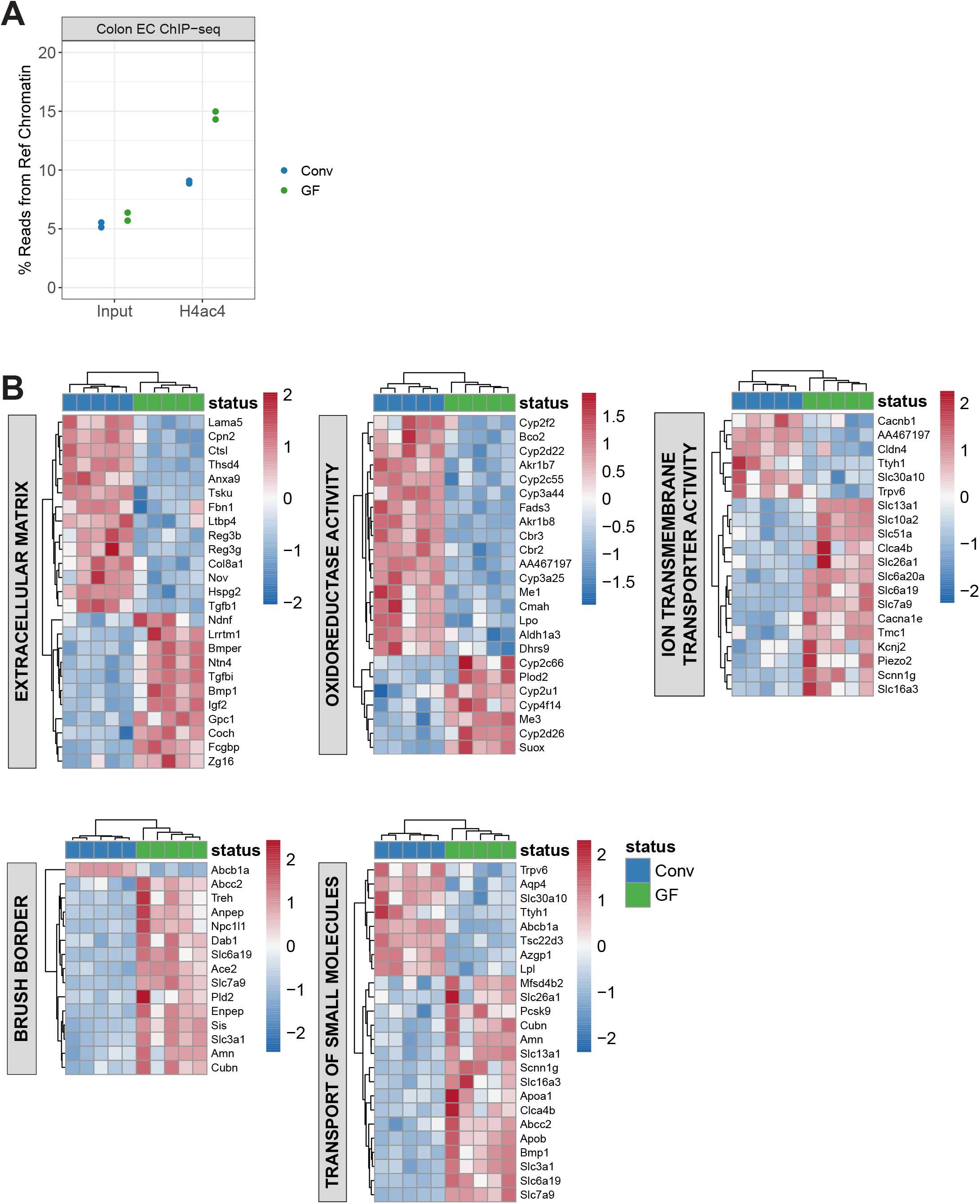
Analysis of H4ac4 ChIP-seq and RNA-seq in colonic epithelial cells from conventional versus germ-free mice. A) The percent of uniquely mapping reads aligning to the *Drosophila* exogenous reference genome is plotted for input and H4ac4 ChIP samples from Conv and GF mice. B) Heatmaps of differentially expressed genes belonging to significantly enriched pathways (GO or Reactome) in colonic epithelial cells from Conv versus GF mice.

**Supplemental Figure 3.**
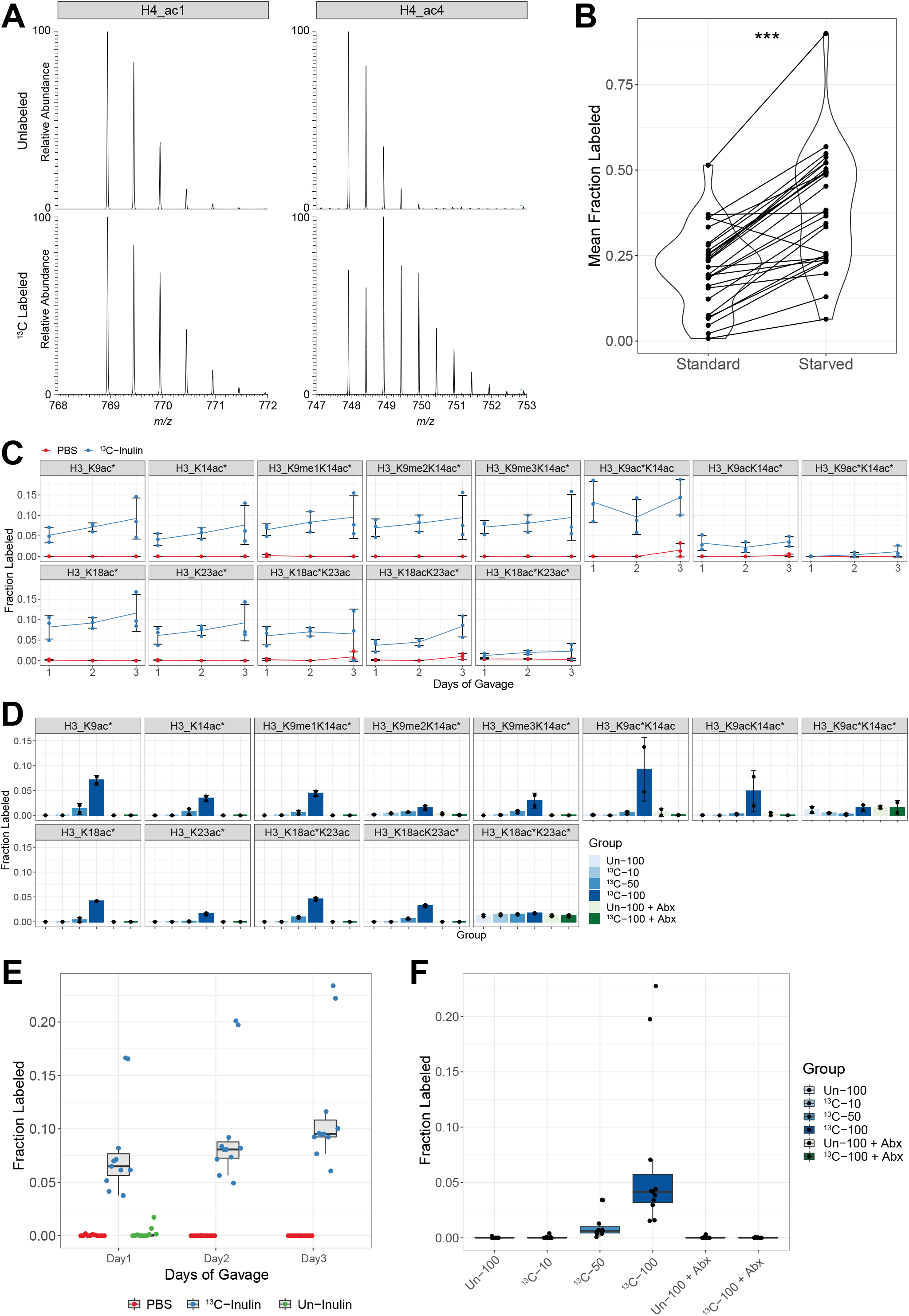
Detection of isotope incorporation into histone acetyl groups after treatment with ^13^C-butyrate in cell culture and ^13^C-inulin in mice. A) Mass spectra for precursor peptide ions corresponding to the histone H4 tail peptide (^4^GKGGKGLGKGGAKR^17^) carrying one (ac1) or four (ac4) acetyl groups after incubation of Caco2 cells with 1 mM unlabeled or U-^13^C-labeled butyrate for 24 hrs. B) Caco2 cells were incubated with 1 mM U-^13^C-butyrate under standard or starved (1% dialyzed FBS – glucose – pyruvate) conditions for 24 hrs. Isotope incorporation into 28 acetylated peptides from histones H3 and H4 is shown with lines connecting the same peptides across conditions. Each point represents the mean of 2 technical replicates. *** p = 2.75 x 10^-9^ by two-tailed, paired t-test. C) Isotope incorporation over time for 13 acetylated histone H3 peptides in colonic epithelial cells from mice receiving daily gavages of ^13^C-inulin or PBS (mean ± sd, n = 3). The top row shows peptides encompassing H3K9ac and H3K14ac while the bottom row shows peptides encompassing H3K18ac and H3K23ac. D) Isotope incorporation for 13 acetylated histone H3 peptides in colonic epithelial cells from mice receiving two daily gavages of labeled (^13^C) or unlabeled (Un) inulin at various doses (10, 50, 100 mg) with or without antibiotic treatment (Abx). Mean ± sd, n = 2. E, F) Summary of isotope incorporation measurements for acetylated histone H3 and H4 peptides from ^13^C-inulin experiments. Each point represents the mean fraction labeled of an acetylated peptide (n = 2-3 mice and 11 peptides). Only mono-acetylated peptides from histones H3 and H4 with one labeled acetyl group are displayed, including H3K9ac*, H3K14ac*, H3K9me1K14ac*, H3K9me2K14ac*, H3K9me3K14ac*, H3K18ac*, H3K23ac*, H4K5ac*, H4K8ac*, H4K12ac*, and H4K16ac*.

**Supplemental Figure 4.**
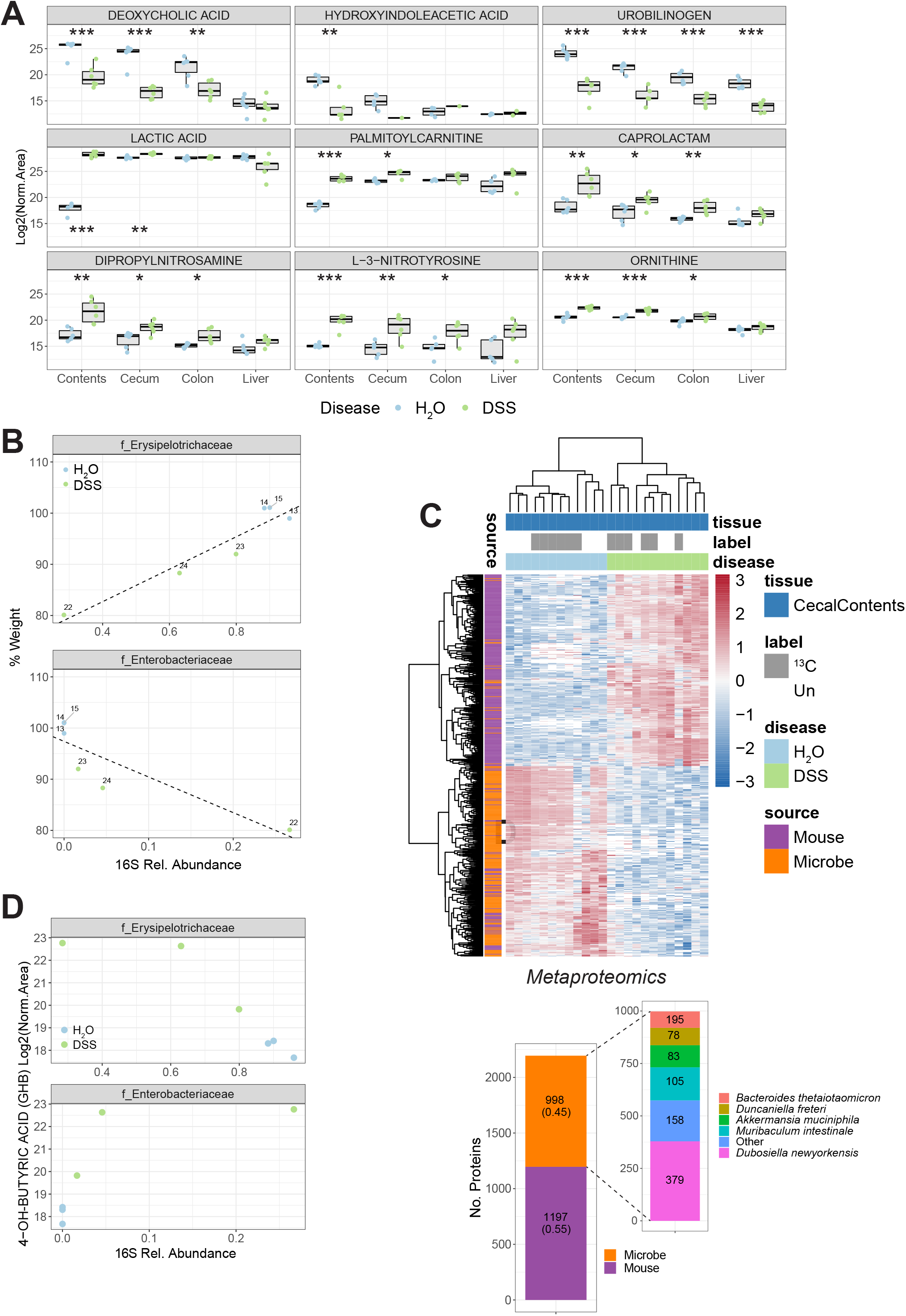
Characterization of differences in metabolites, proteins, and microbiota composition between healthy and colitic mice. A) Normalized abundances of selected metabolites with significant differences. Both groups received Un inulin. Points indicate individual replicates (n = 6). * p < 0.05, ** p < 0.01, *** p < 0.001 by unpaired, two-tailed t-test. B) Relative abundances of different families, representing OTU1 and OTU3, versus relative weight change for each mouse (n = 3 per group). C) Top, heatmap of murine and microbial proteins showing differential abundance in the cecal contents of DSS mice by mass spectrometry (586 features, q < 0.01). Missing values are displayed in white. Below, a summary of protein identifications assigned to mouse versus microbial origin. A more detailed classification of the microbial proteins is depicted in the inset. D) The relative abundances of OTUs from 16S sequencing were correlated (Kendall rank-order method) with the normalized abundances of metabolites in the cecal contents. OTU abundances are plotted against GHB levels for individual mice.

**Supplemental Figure 5.**
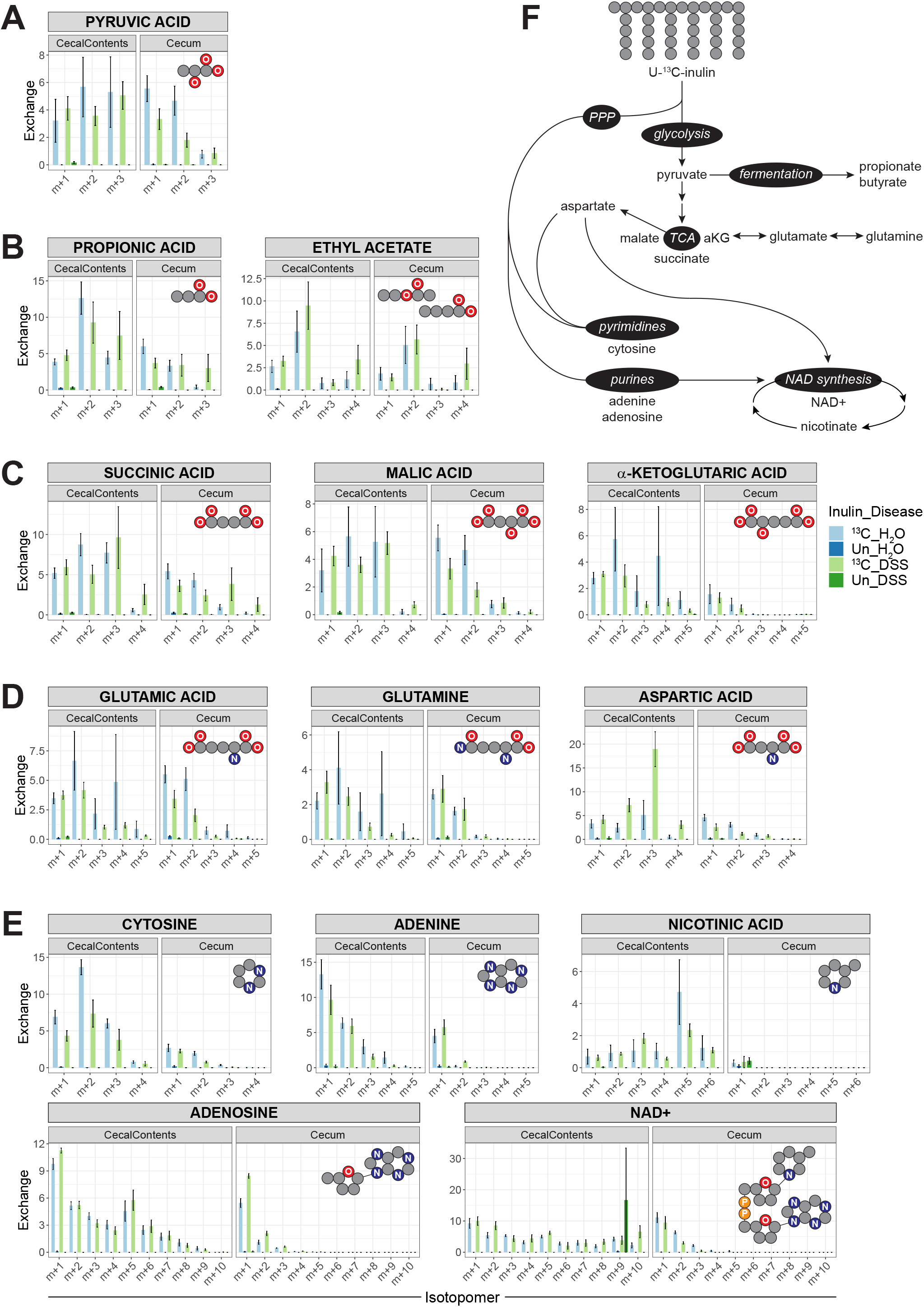
Contribution of dietary fiber to central carbon metabolism, nucleotide metabolism, NAD metabolism, and amino acid metabolism. A-E) Isotopomer distributions for selected metabolites in the cecal contents and cecum from various pathways, including products related to glycolysis (A, pyruvate), fermentation (B, propionate and butyrate/ethyl acetate), TCA cycle (C, succinate, malate, alpha-ketoglutarate), amino acid synthesis (D, glutamate, glutamine, aspartate), and nucleotide and NAD synthesis (E, cytosine, adenine, nicotinate, adenosine, NAD+). In all cases, the m+0 peak is omitted for clarity. For NAD+, isotopomer peaks past m+10 are also omitted. Insets display simplified representations of carbon skeletons. Mean ±sem, n = 6. F) Schematic of putative routes of inulin metabolism.

**Supplemental Figure 6.**
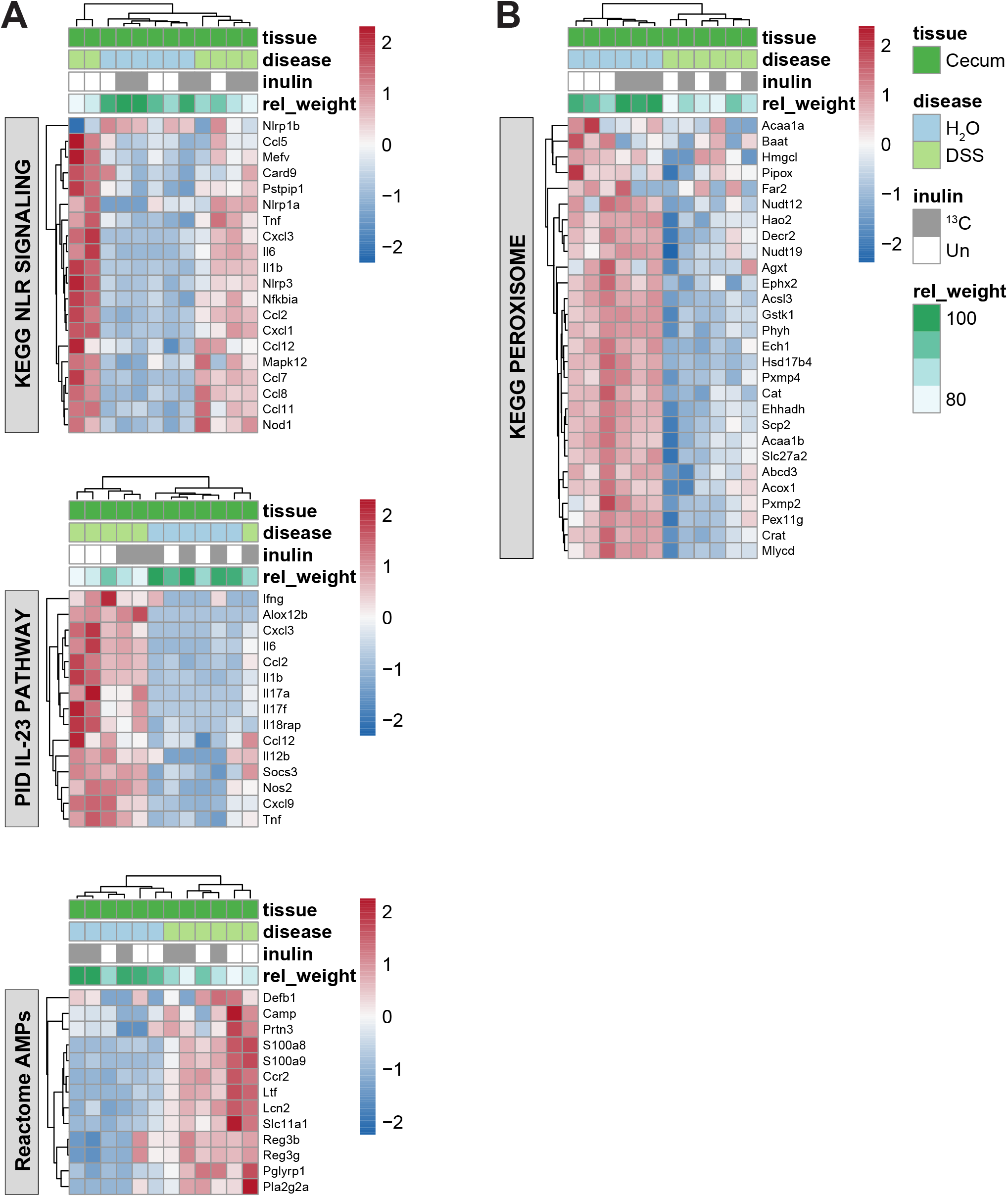
Transcriptional signatures present in the cecal tissue of healthy and colitic mice. A, B) Heatmaps of GSEA core enrichment genes related to pathways upregulated (A) or downregulated (B) in diseased cecal tissue.

**Table 1. Analysis of gene expression in colonic epithelial cells from germ-free versus conventional mice.** This table summarizes results from the RNA-seq performed on colonic epithelial cells from germ-free and conventional mice. The log2 fold-change is calculated as log2(GF/Conv).

**Table 2. Analysis of the overall levels and isotopic labeling of small molecules in tissues from healthy and colitic mice by LC-MS-based untargeted metabolomics.** This table summarizes results from the metabolomics analysis performed on the cecal contents, ceca, colons, and livers of healthy control mice and mice with DSS-induced colitis. This file includes six tabs. The first tab (a) reports qualitative and quantitative information on features across all four LC-MS methods. Normalized peak areas represent peak areas after standardization to a constant extraction concentration of 20 mg/ml. The second tab (b) and third tab (c) report statistics using 2-way ANOVA (disease vs. tissue) and t-tests (DSS vs. H2O within a given tissue) to compare normalized peak areas across conditions. The fourth tab (d) provides a comparison of relative exchange rates in the U-^13^C-inulin and unlabeled conditions to identify ^13^C-labeled molecules within a given tissue and disease state. P-values are from unpaired t-tests. The fifth tab (e) contains the 288 features classified as labeled. The last tab (f) contains data on the isotopomers of these labeled features across all tissues.

**Table 3. Analysis of microbiota composition in healthy and colitic mice by 16S sequencing.** This table details the results from the 16S sequencing performed on the cecal contents of healthy control mice and mice with DSS-induced colitis. Both groups received gavages of ^13^C-inulin. The file includes a tab with the relative abundances of each OTU from each sampled mouse, a tab with a summary of the BLAST searches, and tabs with expanded BLAST results for each OTU.

**Table 4. Metaproteome analysis of the cecal contents from healthy and colitic mice.** This table reports the MSstats results from the mass spectrometry analysis performed on proteins extracted from the cecal contents of healthy control mice and mice with DSS-induced colitis. Both groups received gavages of either unlabeled or U-^13^C-inulin. Log2 fold-changes (DSS/H2O) were calculated separately for these treatments as noted in the Label column (un: unlabeled inulin, 13C: isotope-labeled inulin). Tentative assignments to microbial organisms are based on the 16S sequencing data, which was used to create a metaproteome sequence database for spectrum searching.

**Table 5. Analysis of gene expression in cecum and colon tissue from healthy and colitic mice.** This table summarizes results from the RNA-seq performed on frozen tissue from healthy control mice versus mice with DSS-induced colitis. Both groups received gavages of either unlabeled or U-^13^C-inulin. Within a given tissue and disease state, replicates from the unlabeled and isotope-labeled inulin conditions (n = 3) were considered as one group (n = 6) since the isotope label did not have a strong effect on gene expression. The log2 fold-change is calculated as log2(DSS/H2O). The file includes separate tabs for cecum and colon tissue.

